# A comprehensive single-cell expression atlas of human AML leukemia-initiating cells unravels the contribution of HIF pathway and its therapeutic potential

**DOI:** 10.1101/2022.03.02.482638

**Authors:** Talia Velasco-Hernandez, Juan L. Trincado, Meritxell Vinyoles, Adria Closa, Francisco Gutiérrez-Agüera, Oscar Molina, Virginia C Rodríguez-Cortez, Paolo Petazzi, Sergi Beneyto-Calabuig, Lars Velten, Paola Romecin, Raquel Casquero, Fernando Abollo-Jiménez, Rafael Díaz de la Guardia, Patricia Lorden, Alex Bataller, Helene Lapillonne, Ronald W Stam, Susana Vives, Montserrat Torrebadell, Jose Luis Fuster, Clara Bueno, Eduardo Eyras, Holger Heyn, Pablo Menéndez

## Abstract

Relapse remains a major challenge in the clinical management of acute myeloid leukemia (AML), and is driven by rare therapy-resistant leukemia-initiating stem cells (LSCs) that reside in specific bone marrow niches. Hypoxia signaling keeps cells in a quiescent and metabolically relaxed state, desensitizing them to chemotherapy. This suggests the hypothesis that hypoxia contributes to AML-LSC function and chemoresistance and is a therapeutic target to sensitize AML-LSCs to chemotherapy. Here, we provide a comprehensive single-cell expression atlas (119,000 cells) of AML cells and AML-LSCs in paired diagnostic-relapse samples from risk-stratified patients with AML. The HIF/hypoxia pathway is attenuated in AML-LSCs compared with differentiated AML cells, but is enhanced when compared with healthy hematopoietic cells. Accordingly, chemical inhibition cooperates with standard-of-care chemotherapy to impair leukemogenesis, substantially eliminating AML-LSCs. These findings support the HIF pathway as a stem cell regulator in human AML, and reveal avenues for combinatorial targeted and chemotherapy-based approaches to specifically eliminate AML-LSCs.

## Introduction

Acute Myeloid Leukemia (AML) is the most common acute leukemia in adults, and constitutes a heterogeneous group of disorders characterized by the rapid expansion and accumulation of poorly differentiated myeloid cells in the bone marrow (BM) and infiltrating tissues. Disease heterogeneity is well documented and patients are typically stratified based on cytogenetic, molecular, and immunophenotypic features. While our understanding of the molecular and phenotypic characteristics of AML has substantially improved in recent years, many patients fail to respond to standard-of-care chemotherapy or show early relapse (1, 2).

AML is a paradigm of the hierarchical cancer stem cell model (3). Robust experimental data demonstrate that relapse is mediated by a rare subpopulation of cells, termed leukemia stem cells (LSCs), which are chemotherapy resistant and drive disease recurrence, leading to a more genetically-heterogeneous and clonally-evolved AML (4–6). AML-LSCs share unique properties with normal hematopoietic stem and progenitor cells (HSPCs), including quiescence, resistance to apoptosis and elevated drug efflux, making them partially refractory to chemotherapy.

Hypoxia represents a self-security mechanism to maintain cells in a dormant state, preventing their exhaustion and proliferative damage. Recent data suggest that the rate of oxygen consumption and cell metabolism, rather than oxygen perfusion, is responsible for the hypoxic nature of the BM niche where the LSCs/HSPCs reside (7, 8). Cells respond to hypoxia by activating specific pathways modulated by the hypoxia inducible factors (HIFs), which trigger the expression of hypoxia-regulated genes with key roles in numerous biological processes such as cell proliferation, survival, apoptosis, angiogenesis, metabolism and differentiation (9). At a molecular level, the HIFs constitute a family of three related heterodimeric transcription factors (HIF-1, HIF-2 and HIF-3) whose regulation depends on the oxygen-dependent stabilization of an associated α subunit. Above 5% oxygen, the α subunit is degraded by the proteasome, whereas under hypoxic conditions, it is stabilized post-translationally, dimerizes with the constitutively expressed β subunit and promotes the transcription of HIF target genes (9).

The HIF/hypoxia pathway is important not only for steady-state hematopoiesis, but also for the initiation, progression and chemoresistance of solid tumors and leukemias. Indeed, treatment-resistant AML cells preferentially locate in the hypoxic endosteal niche of the BM, which offers protection from the pro-apoptotic effects of the standard-of-care chemotherapeutic agent cytarabine (AraC) (10). Several studies have shown that, similar to what is observed for normal HSPCs (11), that the loss of HIF-1/2 leads to the complete abrogation of LSCs in different types of human AML and in murine models of chronic myeloid leukemia (12–14), whereas other studies have reported that loss of HIF-1/2 does not impact LSCs in murine models of AML, or can even trigger a more severe leukemic phenotype (15–19). Despite these conflicting observations, HIF-inhibiting drugs are being actively explored as therapeutic agents for AML (12, 13, 20). However, little information is available regarding HIF in human primary AML-LSCs, and importantly, the cytogenetic/molecular heterogeneity intrinsic to AML biology makes it plausible that the action of HIF varies among risk-stratified AML patients.

Here, we used single-cell RNA sequencing (scRNA-seq) to survey the transcriptome of human AML-LSCs in paired diagnostic (Dx) and relapse (REL) samples from risk-stratified patients with AML. Furthermore, we investigated the role of HIF/hypoxia signaling in LSCs quiescence and chemoresistance using cutting-edge *in vitro* and *in vivo* approaches. We found that while the HIF/hypoxia pathway is more weakly expressed in LSCs than in more differentiated AML cells, its chemical inhibition cooperates with chemotherapy to control leukemogenesis, substantially eliminating AML-LSCs. These findings confirm HIF/hypoxia pathway as a stem cell regulator in human AML and offer new avenues for combinatorial targeted and chemotherapy-based approaches to specifically eliminate AML-LSCs.

## Results

### Hypoxia transcriptional signature clusters inv(16) AML subgroup

To capture the contribution of hypoxia pathway in human AML-LSCs, we utilized two publicly available RNA-seq transcriptomic datasets (TARGET (21), including pediatric and adolescent/young patients, and Leucegene (22), including adult patients) generated from Dx samples of the following molecularly defined AML subgroups: (i) inv(16) (CBF-MYH11, n=20/18 patients, TARGET/Leucegene), (ii) t(8;21) (RUNX1- RUNX1T1, n=21/16 patients) and (iii) NPM1^mut^ (n=6/7 patients) as low risk AML; and (iv) MLL-rearranged (MLLr, KMT2A fusions, n=13/15 patients), (v) FLT3^ITD^ (n=4/6 patients), and (vi) normal karyotype (NK, neither chromosomal rearrangements nor NPM1^mut^ or FLT3^ITD^ n=14/10 patients) as intermediate-high risk AML (23). AML samples mutated for *TET2*, *IDH1* or *IDH2* were excluded from analyses because such mutations have been suggested to interfere with HIF signaling (24, 25). A total of 78 samples (147 runs) and 72 samples (301 runs) were analyzed from TARGET and Leucegene, respectively (**Table S1**). For initial data inspection, we used either a multidimensional scaling reduction (MDS) of the genome-wide information or a specific set of 119 hypoxia target genes (Hypoxia signature) characterized by hypoxia-dependent transcriptional induction and by the presence of functional hypoxia response elements (26) (**Table S2**, **Figure 1A**).

**Figure 1.**
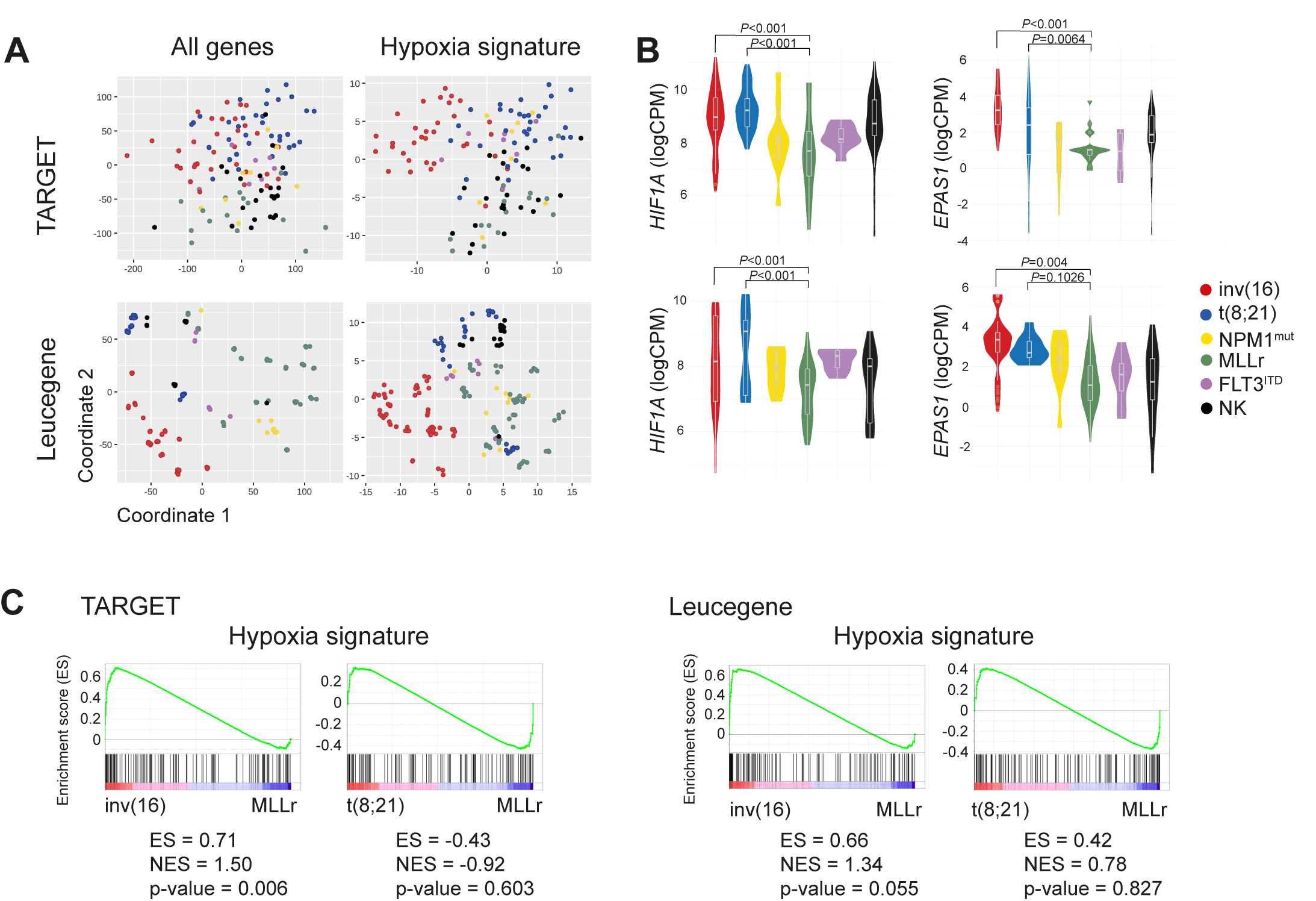
HIF pathway gene expression signature in different AML cytogenetic subgroups. **A.** MDS representation of AML samples from TARGET (78 patient samples and 147 runs) and Leucegene (72 patient samples and 301 runs) databases analyzing the expression of all the detected genes (left panels) or, specifically, the 119 HIF target genes (right panels). **B.** Expression (LogCPM) of *HIF1A* and *HIF2A* (*EPAS1*) in each cytogenetic AML subgroup from TARGET and Leucegene. **C.** GSEA of the HIF pathway comparing inv(16) and t(8;21) with MLLr AMLs. CPM: counts per million; ES: enrichment score; NES: Normalized enrichment score.

The hypoxia signature enabled clustering of the inv(16) AML samples separately from the other cytogenetic groups, in both datasets (**Figure 1A**). The t(8;21) AML samples also clustered partially separately in TARGET and Leucegene datasets, albeit to a lesser extent, whereas MLLr AMLs did so in Leucegene but not in TARGET (**Figure 1A**). The highest expression of *HIF1A* and *HIF2A* (*EPAS1*) was observed in inv(16) and t(8;21) AML samples, both core binding factor (CBF)-rearranged AMLs, whereas MLLr AML samples showed a trend for the lowest expression (**Figure 1B**). Consistently, and in line with data from BloodSpot (27, 28), gene set enrichment analysis (GSEA) revealed a significant enrichment of the hypoxia signature in inv(16), but not in t(8;21) samples, as compared with MLLr samples (**Figure 1C**). Subsequent analyses focused on inv(16) and MLLr subgroups as HIF^high^ and HIF^low^ AML prototypes, respectively. We also included t(8;21) as an additional CBF-rearranged AML, as it has been previously reported to cooperate with HIF1A for leukemogenesis (29).

### Identification of human AML-LSCs using scRNA-seq

To resolve the transcriptional heterogeneity of AML and to survey the expression of HIF pathway genes in AML-LSCs, we performed scRNA-seq on 11 Dx BM samples from pediatric/young adult patients: inv(16) (n=4), t(8;21) (n=3) and MLLr (n=4) (**Figure 2A-B and Figure S1**). Acknowledging the extremely low frequency of LSCs (30, 31), we performed scRNA-seq on two fluorescence activated cell sorting (FACS)- sorted AML populations: CD34+CD38- cells, enriched in LSCs (3, 30, 32) and CD34-CD38+, which are differentiated cells depleted of LSCs (**Figure 2C-D and S1**). A total of 26,976, 19,731 and 24,854 cells were sequenced from inv(16), t(8;21) and MLLr AML subgroups, respectively (**Table S3**). All samples within each cytogenetic-molecular subgroup were computationally integrated and displayed using uniform manifold approximation and projection (UMAP) visualizations (**Figure S2A**). Consistent with the immunophenotype (**Figure S1**), inv(16) and t(8;21) samples expressed high CD34 levels, which was confirmed by scRNA-seq on the sorted population (**Figure 2E and S2A**). By contrast, MLLr samples mainly consisted of CD34- cells, in line with previous studies locating the LSC population within the CD34-CD38+ population in MLLr AML samples (33).

**Figure 2.**
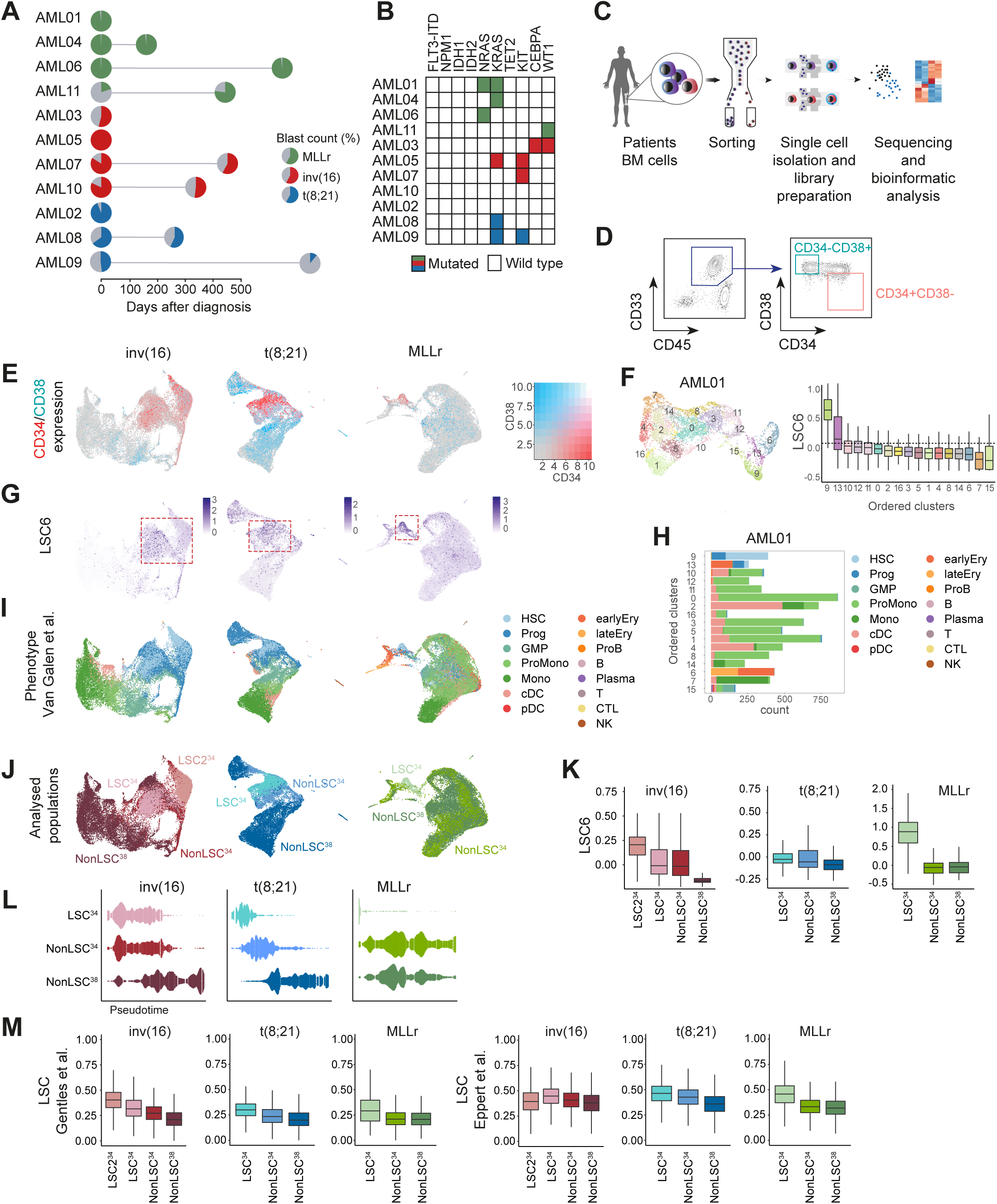
Enrichment and identification of the LSC compartment in the scRNA-seq dataset. **A.** Overview of the primary AML samples used for the scRNA-seq analysis. The distinct cytogenetic subgroups are color-coded. The colored area of the pie-charts depicts the percentage of blasts. Paired- relapsed samples are depicted with a second pie-chart at the time of relapse. Further information of each sample can be found on **Table S3**. **B.** Mutational profile of the analyzed samples. **C.** Scheme depicting the different steps from sample sourcing to scRNA-seq analysis. **D.** Representative FACS profile depicting how the CD34+CD38- and CD34-CD38+ AML cells were FACS- purified for scRNA-seq. The specific FACS profiles of each AML sample can be found in **Figure S1**. **E.** UMAP plots showing the expression of CD34 and CD38 among all cells integrated from different samples in each cytogenetic subgroup. **F.** UMAP plot showing the random clusterization of the cells from the sample AML01 and boxplot of the LSC6 score (Elsayed *et al*) of each cluster for the identification of the LSC-enriched cluster. Dotted line marks the 9^th^ decile. **G.** UMAP plots depicting the LSC6 score assigned to each cell. All cells from the different samples in each cytogenetic subgroup are integrated. Red square marks the LSC6-enriched area. **H.** Number of cells from each predicted phenotype according to Van Galen *et al,* included in each cluster identified in sample AML01. **I-J.** UMAP plots showing the predicted phenotype of the cells according to Van Galen *et al* (**I**), and the assigned population (LSC^34^, NonLSC^34^ and NonLSC^38^) (**J**) for downstream analysis. All cells from the different samples in each cytogenetic subgroup are integrated. **K.** LSC6 (Elsayed *et al*) score of each of the defined populations (LSC^34^, NonLSC^34^ and NonLSC^38^). **L.** Trajectory/Pseudotime analysis of the defined populations from the different cytogenetic subgroups. **M.** Expression of the LSC signatures described by Gentles *et al* and Eppert *et al* in each of the defined populations for the different cytogenetic subgroups. HSC: hematopoietic stem cell; Prog: progenitor; GMP: granulocyte-macrophage progenitor; ProMono: promonocyte; Mono: monocyte; cDC: conventional dendritic cells; pDC: plasmacytoid dendritic cells; Ery: erythroid progenitor; ProB; B cell progenitor; B: mature B cell; Plasma: plasma cell; T: naïve T cell; CTL: cytotoxic T lymphocyte; NK: natural killer cell; LSC: leukemic stem cell; log2FC: log2 fold change.

As the LSC definition relies on functional assays, and CD34 and CD38 are not absolute markers to identify human LSCs, we used the scRNA-seq data to phenotypically categorize *bona fide* LSCs. We performed unsupervised clustering of all cells and utilized the recently published LSC6 score (34), an updated signature of the LSC17 score adapted for pediatric-young AML cell annotations (35). Clusters from each sample were ranked according to LSC6 score values, and only those with the highest LSC6 score were considered enriched in LSCs (**Figure 2F**). When integrating individual samples from the same cytogenetic subgroup, we observed that cells identified as LSCs (highest LSC6 score) clustered together in the integrated UMAPs (**Figure 2G**). Notably, high LSC6-scoring cells colocalized with CD34+CD38- cells across the 3 AML subgroups, including MLLr AML.

We next queried the normal stem/progenitor phenotypic prediction of the LSC6 signature (**Figure 2H-I**). For this, we projected our scRNA-seq data onto an existing reference annotation dataset containing 15 healthy hematologic cell types (36). Each AML sample was first projected individually (**Figure 2H** and https://github.com/JLTrincado/scAML) and then all AML samples within the same cytogenetic subgroup were integrated (**Figure 2I**). We observed that phenotypically identical annotated clusters colocalized together in the UMAP, demonstrating a similar identity/phenotype across different patients. Total CD34+ cells and high LSC6-scoring cells (identified as LSCs) were enriched for hematopoietic stem cells (HSCs) and progenitors (**Figure 2I**). Data projection on an additional annotated dataset (37) confirmed the stemness phenotype of these cells, which overlapped with HSCs, multipotent progenitors (MPPs), lympho- myeloid primed progenitors (LMPPs) and myeloblasts (**Figure S2B-C**).

This LSC-enriched CD34+CD38- cluster (hereinafter, LSC^34^) was further characterized and compared with the remaining CD34+CD38- cells not identified as LSCs (hereinafter, NonLSC^34^) and with the CD34-CD38+ cells (hereinafter, NonLSC^38^) (**Figure 2J**). Of note, in an individual inv(16) sample (AML10), an additional LSC6^high^ cluster was identified and classified as HSC/progenitors but with high expression of *HBB* (LSC2^34^) (**Figure S2D**). When the different Dx-AML samples were integrated, we consistently observed a lower LSC6 score from the LSC^34^ towards the more differentiated NonLSC^38^ (**Figure 2K**), in accord with the observed pseudotime trajectories of these populations along a continuum of differentiation from LSC^34^ to NonLSC^38^ (**Figure 2L**) irrespective of the AML cytogenetic subgroup. We further explored the relationship between the *in silico* predicted cellular annotations by obtaining their latent space in each molecular subgroup, finding a similar continuum of differentiation from the most undifferentiated cells to terminally differentiated monocytes (**Figure S2E-F**). These analyses not only validated our predictions but also highlight the cellular heterogeneity and diversity of both CD34+CD38- and CD34-CD38+ AML cells. In this line, two additional widely used LSC signatures (30, 38) correlated with the LSC^34^ clusters identified based on the LSC6 signature and stemness projection (**Figure 2M**).

Additionally, we analyzed the expression of 18 specific markers commonly used to identify LSCs in human AML (39–44) and found a panel to be consistently overexpressed in the LSC^34^ cluster identified in inv(16) samples (*CD99*, *CD82*, *CD52*, *CD47*, *IL3RA*), t(8;21) samples (*CD99*, *CD52* and *CD96*) and in MLLr samples (*CD99*, *CD82*, *CD52* and *CD47*), as compared with both NonLSC^34^ and NonLSC^38^ clusters (**Figure S2G**). Finally, to rule out bias in the gene expression analysis due to contaminating healthy HSCs/progenitors with an immunophenotype overlapping that of AML-LSCs (30, 45), the expression of specific genes reported to be upregulated in AML cells was compared against healthy BM obtained from the Human Cell Atlas (46). Results showed that *CLEC12A* (*CLL-1*) (47) and *JUND* were overexpressed in AML cells across cytogenetic groups (**Figure 2SH**), whereas *SPARC,* or *RUNX1T1* and *POU4F1,* or *HOXA9*, *HOXA10* and *PBX3* were specifically upregulated in inv(16), t(8;21), or MLLr AML cells, respectively (**Figure S2H**).

### Transcriptional characterization identifies key molecular features of the AML-LSCs

Recent studies have revealed the existence of dormant and active HSCs in mice (48–51) and humans (52), while AML-LSCs are documented to be quiescent/dormant. To characterize the transcriptional heterogeneity of human AML-LSCs, we first analyzed the cell cycle and quiescence/metabolic dormancy of the LSC^34^ clusters across the cytogenetic groups (**Figure 3A-B**). We took advantage of validated signatures defining the G_0_ cell cycle status (*Neg G0 to G1* [GO:0070317] and G_0_M^high^ (49)) (**Figure 3B and Table S2**). LSC^34^ were consistently found in G_0_/G_1_ cell cycle phase (**Figure 3A**), and the *Neg G0 to G1* and G_0_M^high^ dormancy signatures were enriched in LSC^34^ clusters across the distinct AML molecular subgroups (**Figure 3B**), revealing homogeneous LSC^34^ clusters based on the G_0_ phase and/or quiescence status of the cells.

**Figure 3.**
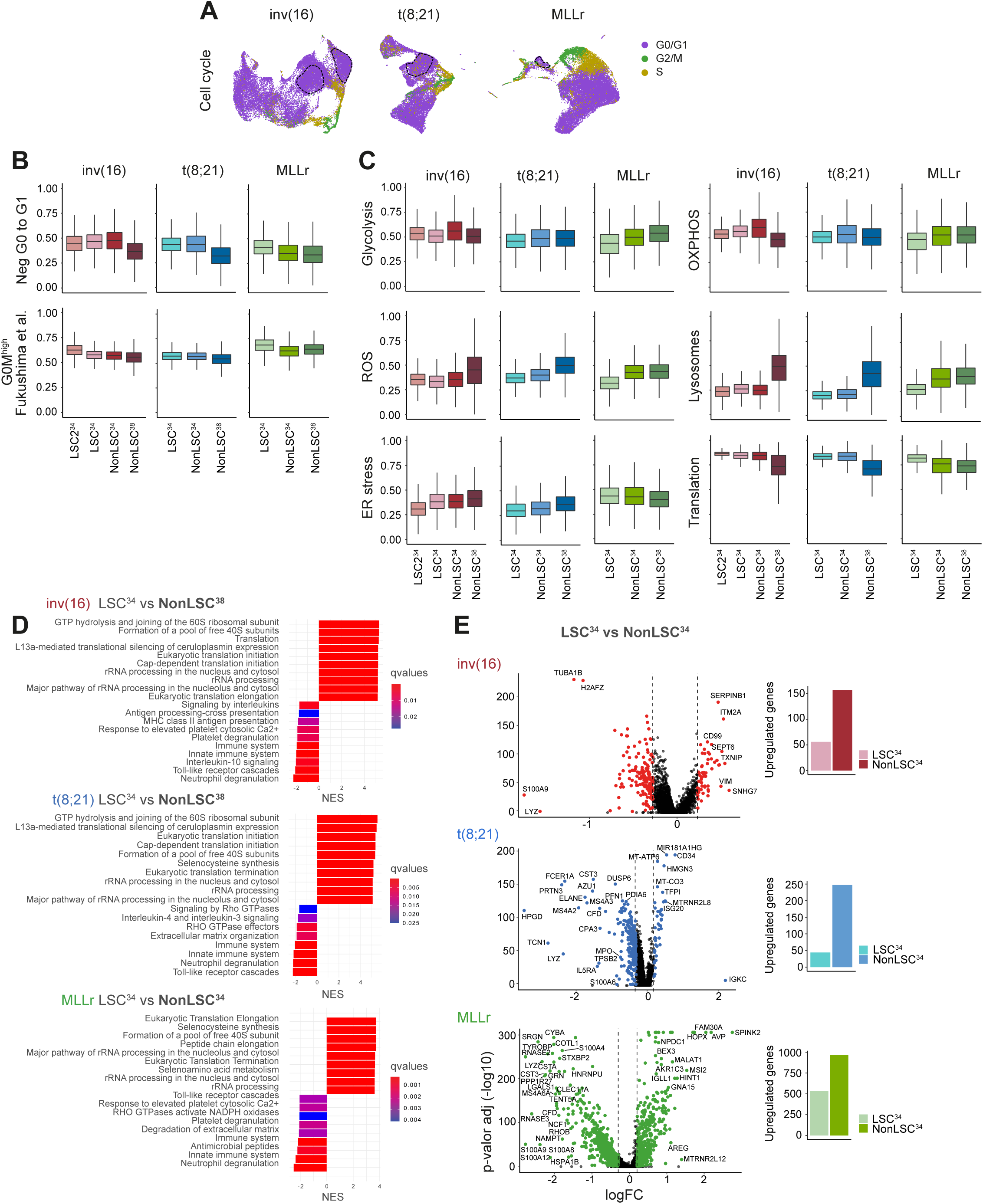
Cell cycle and metabolic characterization of the LSC^34^ cluster. **A.** UMAP plots showing the cell cycle phases prediction for each cell. Cells from all the different samples in each cytogenetic subgroup are integrated. **B.** Quiescence status analysis of the defined populations from the different cytogenetic subgroups using the GO signature *Neg G0 to G1* (GO:0070317) and the dormancy signature G_0_M^high^ described in Fukushima *et al*. **C.** Analysis of different metabolic pathways related to stemness and hypoxia (Glycolysis, OXPHOS, ROS, Lysosomes, ER stress and Translation) for the defined populations from the different cytogenetic subgroups. **D.** GSEA showing the enriched biological pathways in the indicated populations of cells. For inv(16) and t(8;21) AMLs, LSC^34^ cells are compared with NonLSC^38^ cells. For MLLr AML, LSC^34^ cells are compared with NonLSC^34^ cells. Complementary analyses are shown in **Figure S3**. **E.** Volcano plots showing the DEGs between LSC^34^ and NonLSC^34^ cells of each cytogenetic subgroup. Plots in the right show the total number of upregulated genes in each population.

We next analyzed the expression of different metabolic signatures previously related to both HSCs/LSCs and to hypoxia signaling (**Table S2**). Glycolysis (42) signature was less represented in the LSC^34^ in the MLLr AML cells, similarly to oxidative phosphorylation (OXPHOS) (53) (**Figure 3C**). However, OXPHOS was increased in LSC^34^ cells respect to NonLSC^38^ in inv(16) AML cells. Reactive oxygen species (ROS) (42) and lysosome (54) signatures were less represented in the LSC^34^ cluster across the distinct AML molecular subgroups, consistent with lower ROS levels reported in HSCs/LSCs (**Figure 3C**). By contrast, LSC^34^ cells displayed an enrichment in Translation signature consistent with recent publications indicating a high protein production rate in these cells (55–57). However, the ER stress signature (12) differed between LSC^34^ cells from distinct molecular subgroups, being less represented in CBF-rearranged AMLs and enriched in MLLr AMLs (**Figure 3C**).

Unsupervised hierarchical clustering of the differentially expressed genes (DEGs) revealed that in CBF- rearranged AMLs, the LSC^34^ cluster is transcriptionally closer to the NonLSC^34^ cluster than to the NonLSC^38^ cluster (**Figure S3A**). Functional enrichment analysis revealed that the main altered functions between LSC^34^ and NonLSC^38^ clusters were associated with *Translation* and other *Ribosomal-related processes* (**Figure 3D**), whereas functional terms related to *Mitosis* and *Cell cycle* were the most altered between NonLSC^34^ and NonLSC^38^ clusters (**Figure S3B**), confirming that the NonLSC^34^ cluster is more proliferative than the LSC^34^ cluster (**Figure S3B**). In contrast, the LSC^34^ cells in MLLr AML differed transcriptionally from both NonLSC^34^ and NonLSC^38^ cells, which were transcriptionally closer together (**Figure S3A**), in line with the pseudotime analysis (**Figure 2L**). Functional enrichment analysis of MLLr AML samples revealed that the main altered functions between LSC^34^ and both NonLSC^34^ and NonLSC^38^ clusters were associated to *Translation* and *Ribosomal-related processes* (**Figure 3E** and **S3C**). Notably, specific transcriptional features were associated with the LSC^34^ cluster in each molecular AML subgroup. Overall, a higher number of DEGs were upregulated in the NonLSC^34^ cluster with respect to the LSC^34^ cluster (**Figure 3E**, right plots) regardless of the cytogenetic group, suggesting a greater transcriptionally activity in line with the enrichment of the LSC^34^ cluster in dormancy and the G_0_ signature. In total, 56, 44 and 573 genes were found upregulated in the LSC^34^ cluster in inv(16), t(8;21) and MLLr AMLs, respectively (**Figure S4A**). Of these, ten genes were consistently upregulated in the LSC^34^ cluster across all molecular subgroups (*AKR1C3, CD34, CD52, HIST1H2AC, ITM2A, LIMS1, MTRNR2L8, PNISR, SEPT6, SERPINB1*) (**Figure S4B**).

### Low expression of the hypoxia signaling signature in human AML-LSCs

Having captured the transcriptional identity of the LSC^34^ cluster across the three AML molecular subgroups, we sought to analyze the hypoxia signaling pathway in LSC^34^ cells using the aforementioned panel of 119 hypoxia target genes (Hypoxia signature) (**Table S2**) to determine the hypoxia enrichment score. The LSC^34^ cluster consistently showed the lowest hypoxia score across all three cytogenetic subgroups (**Figure 4A- B**), in line with the lowest expression of *HIF1A* (**Figure 4C**). To rule out potential bias in the selection of the 119 genes defining the hypoxia signature, we further employed five transcriptional signatures containing genes upregulated under hypoxia conditions (**Figure 4D** and **Table S2**) and confirmed a uniformly lower hypoxia score in the LSC^34^ cluster with a transition towards enrichment in the hypoxia signature in NonLSC^38^ differentiated AML cells. The poor hypoxia signaling observed in human AML-LSCs was accord with a weaker ROS signature in the LSC^34^ cells (**Figure 3C**). Notably, while the hypoxia signature showed the lowest enrichment score in LSC^34^ among the distinct analyzed clusters, it was routinely enriched in both total AML cells and LSC^34^ cells as compared with both whole healthy BM cells and healthy CD34+ cells, respectively, regardless of the cytogenetic subgroup (**Figure 4E**).

**Figure 4.**
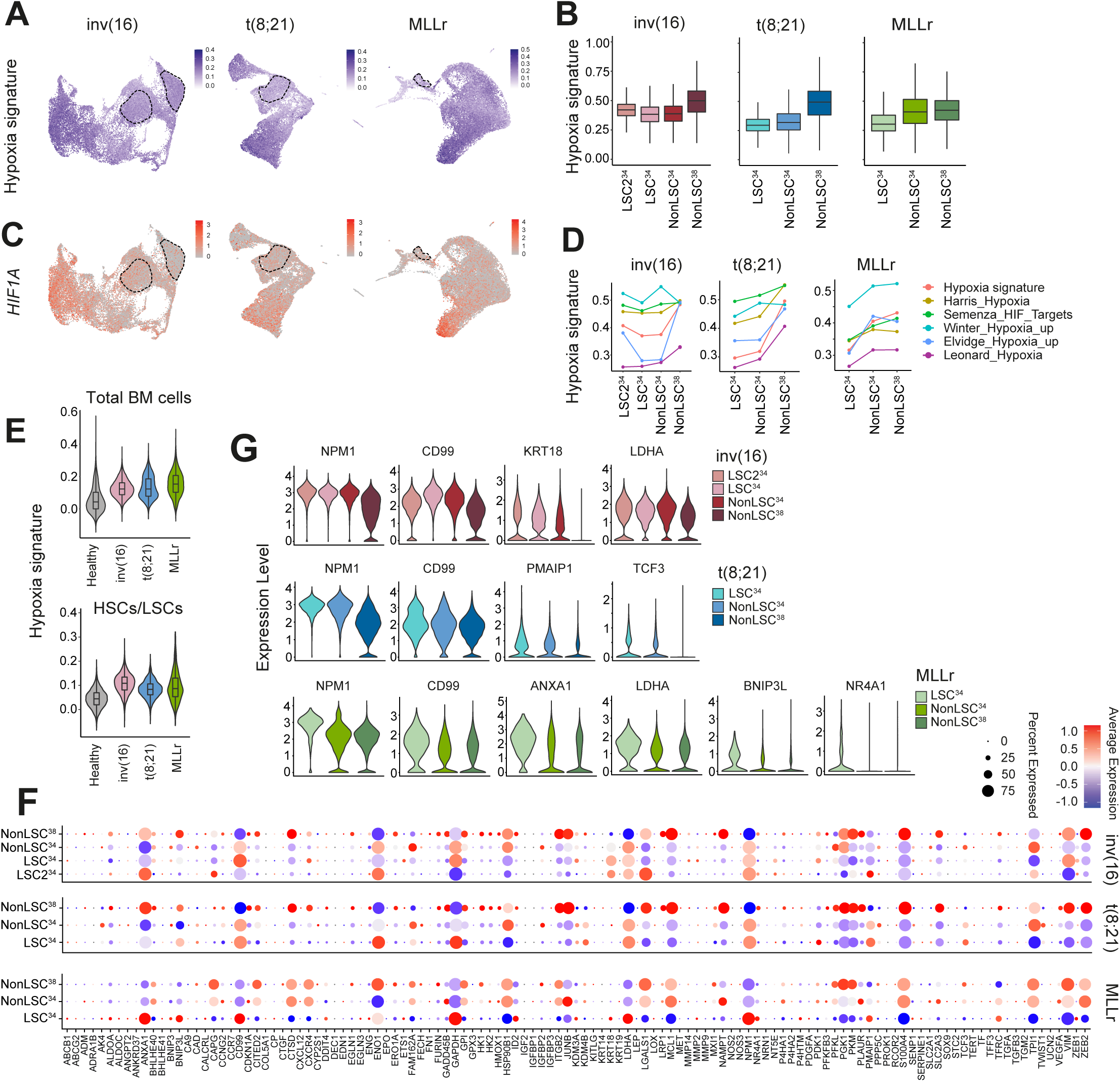
Low expression of hypoxia signaling signature in human AML-LSCs. **A.** UMAP plots showing expression of the hypoxia signature in all cells integrated from the different samples in each cytogenetic subgroup. **B.** Hypoxia signature score in each of the defined populations from the different cytogenetic subgroups. **C.** UMAP plots showing expression of the *HIF1A* gene in all cells integrated from the different samples in each cytogenetic subgroup. **D.** Hypoxia signature score of each of the defined clusters comparing the hypoxia signature used in this study with 5 hypoxia signatures previously reported. **E.** Hypoxia expression signature comparing each AML cytogenetic subgroup with healthy total BM cells (upper plot) or healthy HSCs/LSCs (bottom plot). **F.** Expression of the 119 genes from the hypoxia signature in each of the defined clusters. **G.** Violin plots showing the expression of the significantly overexpressed genes of the hypoxia signaling pathway in the LSC^34^ cluster in each cytogenetic AML subgroup.

Most of the HIF1A targets were upregulated in the differentiated NonLSC^38^ cluster (**Figure 4F**). However, when HIF1A target genes differentially expressed among the three clusters were analyzed in more detail, several HIF1A targets were significantly upregulated in the LSC^34^ cluster: *NPM1*, *CD99, KRT18 and LDHA* in inv(16) samples; *NPM1, CD99, PMAIP1* and *TCF3* in t(8;21) samples; and *NPM1, CD99, ANXA1, LDHA, BNIP3L* and *NR4A1* in MLLr samples (**Figure 4F-G**). Together, although AML-LSCs display a weak hypoxic signature across all the AML subgroups, specific hypoxia-related genes were up-regulated in LSC^34^ cells. Notably, the hypoxia signature was overexpressed throughout different tumoral populations compared with healthy hematopoietic BM cells.

### Paired Dx-REL analysis reveals patient-specific differential molecular features of the AML-LSCs

Chemoresistant LSCs display biological features that differ from those of “therapy naïve” LSCs, including a more diverse phenotype, gene expression changes and an increased metabolic flexibility (4, 5, 58–60). To study the evolution of chemoresistant LSCs underlying AML relapse, we performed scRNA-seq in paired patient-matched Dx-REL samples (**Figure 2A**). In total, 12,005, 15,909 and 19,506 cells were sequenced from inv(16), t(8;21) and MLLr REL-AML patients, respectively. The LSC^34^ cluster was identified separately at Dx and REL before data integration (**Figure 5A** and **S5A**). Results showed a higher transcriptional heterogeneity in REL than in Dx samples, as evidenced by numerous small clusters of cells with a lymphoid or erythroid phenotype (**Figure 5A** bottom plots and **S5B**). Of note, the degree of transcriptional overlap between Dx-REL pairs varied from patient to patient when the total number of cells was integrated (**Figure 5A** and **S5A,** https://github.com/JLTrincado/scAML), suggesting patient-specific transcriptional changes in Dx-REL pairs. Similarly, analysis of the LSC6 score in paired Dx-REL samples also revealed patient-specific heterogeneity with a trend towards an increased LSC6 score at REL (4/7) (**Figure 5B-C** and **S5C**). In addition, analysis of the hypoxia score in paired Dx-REL samples also revealed patient-specific heterogeneity, with an inverse Dx-to-REL evolution with respect to the LSC6 score (6/7) (**Figure 5B-C** and **S5D**). Finally, dormancy, ER stress and ROS signatures also revealed a variable, patient-specific evolution from Dx to REL in LSC^34^ cells irrespective of the AML subgroup (**Figure 5D-E** and **S5E**). Notably, the DEGs found in the Dx-LSC^34^ cluster that were upregulated in the REL-LSC^34^ cluster varied between inv(16), t(8;21) and MLLr AMLs, highlighting molecular subgroup-specific differences (**Figure S5F**). *SERPINB1, PNISR, ITM2A, CD34* and *AKR1C3* were the genes shared across the inv(16) and t(8;21) subgroups.

**Figure 5.**
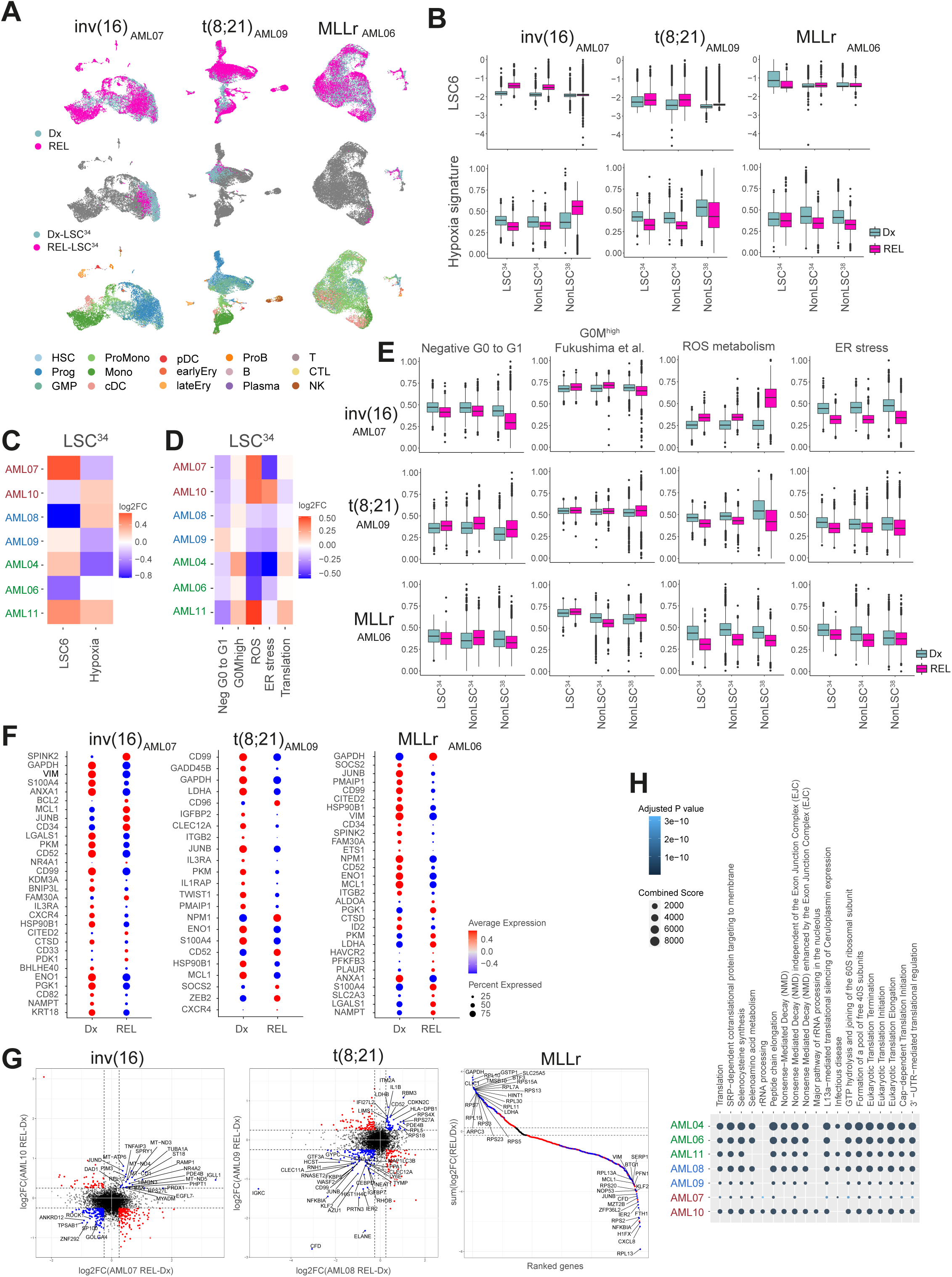
REL-LSC^34^ cluster reveals patient-specific differential molecular features. **A.** UMAP plots integrating patient-matched AML cells at Dx and REL (top plots), showing the identified LSC^34^ cluster at Dx and REL (middle plots) and showing the predicted phenotype according to Van Galen *et al* (bottom plots). One pair from each cytogenetic subgroup is shown. Additional paired-samples are analyzed in **Figure S5A-B**. **B.** LSC6 score (top plots) and hypoxia signature score (bottom plots) of the defined clusters at Dx and REL for each AML cytogenetic subgroup. **C-D.** Clustered representation of the variation of the LSC6 and hypoxia (**C**) and metabolic pathways (**D**) signature scores in the LSC^34^ population in the 7 Dx-REL pairs. **E.** Score of indicated metabolic pathways related to stemness and hypoxia in the defined clusters at Dx and REL for each AML cytogenetic subgroup. **F.** HIF target genes differentially expressed in the LSC^34^ population at Dx *versus* REL in each pair from the indicated patients. Additional paired-samples are analysed in **Figure S5G**. **G.** Comparison of the DEGs in the LSC^34^ population of each paired sample in each cytogenetic subgroup. For inv(16) and t(8;21) AMLs, plots compare 2 AML Dx-REL pairs (AML07 and AML10 for inv(16); AML08 and AML09 for t(8;21)). For MLLr AMLs, plot compares 3 AML Dx-REL pairs (AML04, AML06 and AML11). Blue and red dots depict genes with similar or different, respectively, expression in paired Dx *versus* REL samples. **H.** Reactome showing biological pathways enriched in REL-LSC^34^ cells compared to Dx-LSC^34^ cells. HSC: hematopoietic stem cell; Prog: progenitor; GMP: granulocyte-macrophage progenitor; ProMono: promonocyte; Mono: monocyte; cDC: conventional dendritic cells; pDC: plasmacytoid dendritic cells; Ery: erythroid progenitor; ProB; B cell progenitor; B: mature B cell; Plasma: plasma cell; T: naïve T cell; CTL: cytotoxic T lymphocyte; NK: natural killer cell; LSC: leukemic stem cell; log2FC: log2 fold change.

Specifically, several HIF1A target genes were found differentially expressed in Dx- and REL-LSC^34^ cells (**Figure 5F** and **S5G**). In LSC^34^ from inv(16) AMLs, *HSP90B1* was cosistently down-regulated between Dx and REL in both patients. In LSC^34^ cells from t(8;21) AMLs, four genes (*CD99, JUNB, CLEC12A* and *PMAIP1*) showed a consistent down-regulation between Dx and REL in both patients. Finally, in LSC^34^ cells from MLLr AML samples, six genes showed a consistent change (down-regulation: *JUNB, MCL1* and *VIM*; up-regulation: *GAPDH*, *LDHA* and *PKM*) between Dx and REL in all three patients. In addition to the hypoxia pathway, we analyzed those DEGs showing a consistent change (up- or down-regulation) between Dx and REL in the LSC^34^ cluster for each molecular subgroup (**Figure 5G**), which identified *EGFL7*, *CD52* as well as many ribosomal proteins consistently upregulated in REL samples. Functional enrichment analysis using these genes revealed *Translation*-related terms as the main altered functions in REL-LSC^34^ cells (**Figure 5H**).

### Inhibition of HIF pathway sensitizes AML-LSCs to chemotherapy *in vitro*

HIF signaling and hypoxic BM niches are reported to protect leukemic cells from chemotherapy by promoting quiescence and low metabolic activity (61–63). We found that the HIF pathway signature was less enriched in AML-LSCs than in more differentiated AML blasts, and the hypoxia score in paired Dx-REL samples showed patient-specific heterogeneity. By contrast, the HIF signature was consistently enriched in AML-LSCs as compared with healthy BM cells and HSCs, prompting us to explore its potential therapeutic role. For this, we combined the chemical inhibitor BAY87-2243 (BAY87), which inhibits both HIF1A and HIF2A by preventing their protein accumulation under hypoxia (64), with AraC, a standard-of-care chemotherapeutic in AML (60, 65). The cell lines THP-1 (MLLr), Kasumi-1 (t(8;21)) and ME-1(inv(16)) were treated for 48 h in hypoxic conditions (5% O_2_) with AraC, BAY87 or the combination (combo), and the clonogenic capacity of the resistant cells was assessed by colony-forming unit (CFU) assays (**Figure 6A**). Although response to BAY87 was cell-line dependent, we found an additive effect with AraC in ME-1 cells and a dramatic inhibitory effect of BAY87 (alone or combined with AraC) in THP-1 cells (**Figure 6A**).

**Figure 6.**
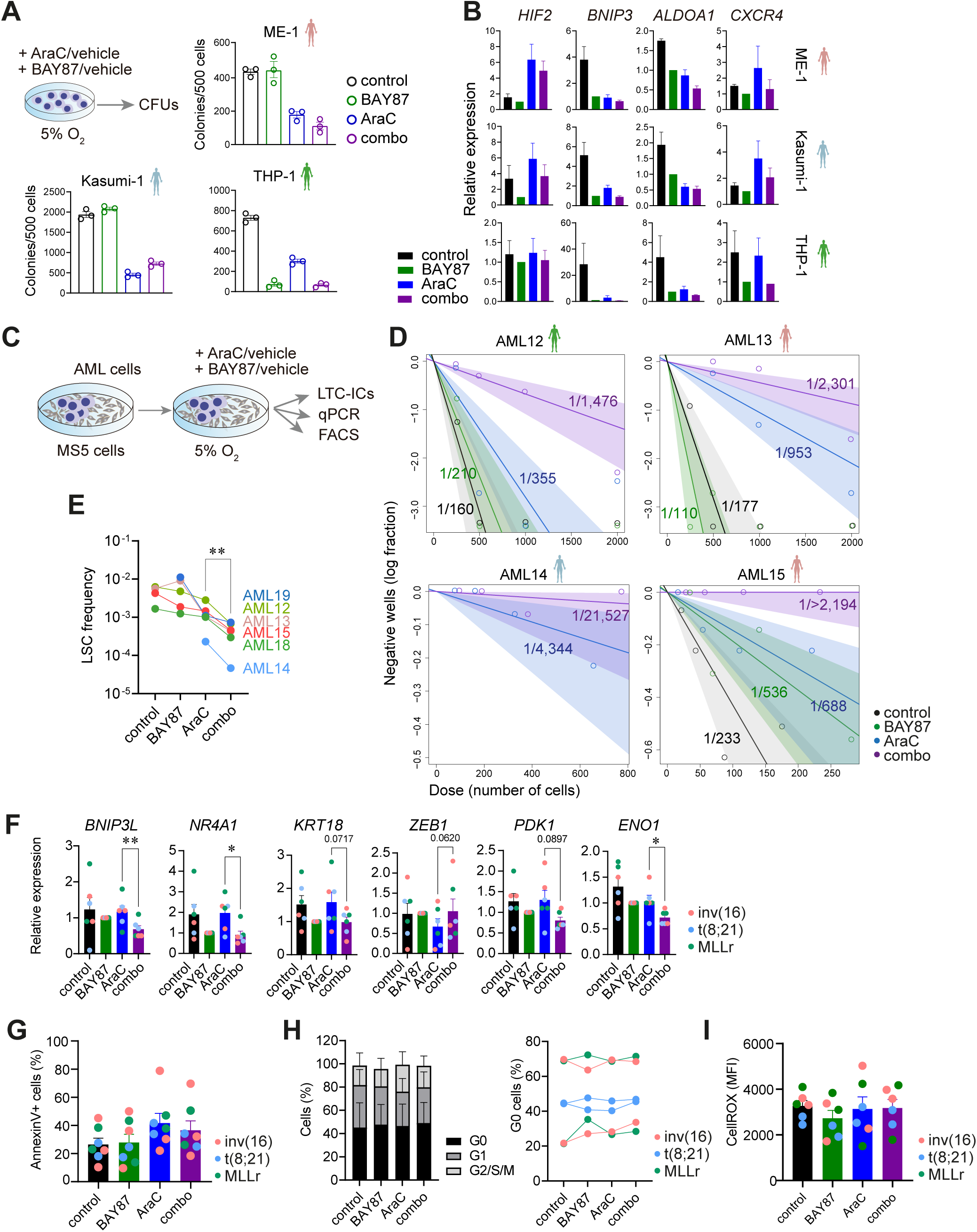
Inhibition of HIF pathway sensitizes AML-LSCs to chemotherapy *in vitro*. **A.** CFU-assays from AML cell lines treated during 48 h with the indicated drugs at 5% O_2_. Results shown from one representative experiment (n=3 technical replicates). **B.** Expression by qPCR of the indicated HIF target genes after 48 h treatment with the indicated drugs at 5% O_2_ (n=3 independent experiments). Expression is normalized respect to the BAY87 samples **C.** Experimental overview for D-I. Human AML primary cells were cultured over MS5 cells for 4 days and treated afterwards with the indicated drugs for 48 h at 5% O_2_. At the completion of the treatment, cells were used for gene expression, flow cytometry or LTC-IC assays (n=15 wells/ treatment and AML sample). **D.** Estimation of the LSC frequency at the completion of the LTC-IC assay calculated using the ELDA software. **E.** Impact of the indicated treatment on the LSC frequency for all the analyzed samples (n=6). Statistical significance was calculated using the Ratio paired Students’t test. *P*-values are indicated for AraC-combo groups comparison. **F.** Expression of the indicated HIF target genes identified in the scRNA-seq analysis to be overexpressed in the LSC cluster after 48 h treatment with the indicated drugs at 5% O_2_ (n=6 samples, AML03, AML16- AML20, 2 per cytogenetic group). Statistical significance was calculated using the paired Students’ t test. Expression is normalized respect to the BAY87 samples. **G-I.** Apoptosis quantification with Annexin V staining (**G**), cell cycle analysis by FACS (**H**) and ROS content measured using CellROX staining (**I**), in AML cells treated with the indicated drugs for 48 h at 5% O_2_ (n=6 samples, AML03, AML16-AML20). Data are shown as mean ± SEM. * *P*<0.1, \*\**P*<0.01.

Quantitative PCR analysis confirmed a decrease in the expression of master HIF1A target genes (*HIF2A, BNIP3, ALDOA1* and *CXCR4)* across the AML cells treated with BAY87 (**Figure 6B**).

We next performed long-term culture-initiating cells (LTC-IC) assays to assess the impact of HIF inhibition on AML-LSCs. Primary cells from six AML patients representing the three cytogenetic subgroups were treated for 48 h in hypoxic conditions (5% O_2_) with AraC, BAY87 or the combo, and a significant decrease in AML-LSC frequency was consistently observed upon treatment with the combo (**Figure 6C-E** and **S6A**). We also analyzed the expression of genes from the HIF pathway identified in our scRNA-seq analysis as differentially expressed in the LSC^34^ compartment in the AML cells after 48 h treatment (**Figure 6F** and **S6B**). BAY87-treated cells showed a decrease in the expression of master genes related to glycolysis (*ENO1* and *PDK1*) and apoptosis (*BNIP3L* and *NR4A1*). We also found a decrease in the expression of *KRT18* related to tumorigenesis and an increase in *ZEB1* expression, in line with its role as a stemness and tumour repressor in AML (66). The presence of chromosomal abnormalities (inv(16), t(8;21) and MLLr) was detected by FISH and/or qPCR at the end of treatments, confirming that LTC-ICs originated from the original leukemic clone and not from residual healthy myeloid progenitors (**Figure S6C-D**). Of note, addition of BAY87 to AraC treatment did not impact apoptosis, cell cycle status or ROS content in the therapy-resistant AML cells (**Figure 6 G-I**). Overall, these data suggest that HIF inhibition may sensitize bulk AML cells and, more importantly, AML-LSCs, to AraC-based standard-of-care treatment, independently of the AML cytogenetic subgroup.

### Inhibition of HIF pathway sensitizes AML-LSCs to chemotherapy *in vivo*

We next aimed to address the impact of HIF inhibition alone or combined with AraC on AML-LSCs *in vivo* (**Figure 7A**). Because low-risk CBF-rearranged (inv(16) and t(8;21)) AMLs have been extensively reported to be very inefficient in engrafting immunodeficient mice (6, 67), we focused our *in vivo* studies on MLLr AMLs. Immunodeficient (NSG) mice were intra-BM-transplanted with primary MLLr AML cells and mice were randomized into the following treatment groups when AML graft levels were detectable in BM: (i) control, (ii) AraC, (iii) BAY87 and (iv) combo (**Figure 7B**). Primografts were treated for five days and mice were sacrificed and analyzed 72 h later (day 8), ensuring clearance of AraC and its metabolites, as previously reported (60, 68). Compared with control mice, peripheral cytopenias (leucopenia, anemia and trombocytopenia) (**Figure S7A**) and a decreased percentage (**Figure S7B**) and total number (**Figure 7C**) of live cells in BM were observed in AraC-treated mice, confirming the cytoreductive/cytostatic effect of the treatment. Notably, BAY87 synergized with AraC to reduce the leukemic burden in peripheral blood (PB), BM, spleen and liver (**Figure 7D** and **S7C**). The clonogenic and stemness potential of the treated primograft cells were next assessed *ex vivo* in CFU-assays. Primograft AML cells from combo-treated mice showed 2-fold less clonogenic potential than counterparts from AraC-treated mice (**Figure 7E**, left panel). In addition, the resulting colonies from combo-treated primograft AML cells were much smaller and with 4-fold less cellularity than those from AraC-treated primograft AML cells (**Figure 7E**, middle panel). The presence of the MLLr was detected by FISH in cells collected from the CFUs, confirming colonies originated from the transplanted MLLr leukemic cells and not from residual healthy myeloid progenitors (**Figure 7E**, right panel).

**Figure 7.**
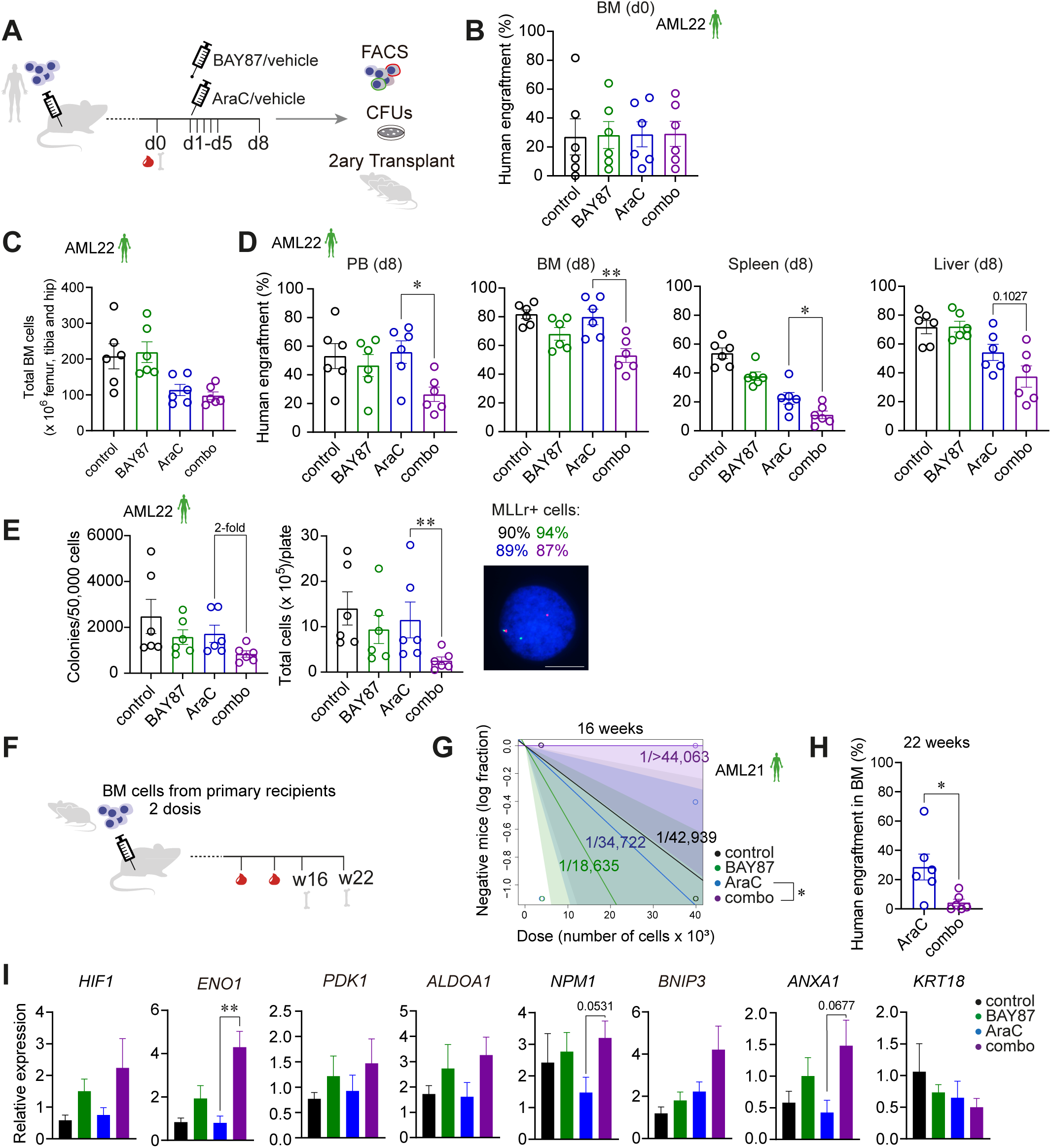
Inhibition of HIF pathway sensitizes AML-LSCs to chemotherapy *in vivo*. **A.** Scheme of the experimental design. Human AML-engrafted mice were treated with the indicated drugs for 5 days. After completion of the treatment, organs were collected and analyzed by FACS. Cells from the BM were used for *ex vivo* CFU assays and secondary transplantations. **B.** Representative human engraftment in BM before treatment (n=6 mice/group) (n=3 independent experiments). **C.** Representative total live BM cells (mouse and human) in mice after each indicated treatment (n=6/group) (n=3 independent experiments). **D.** Representative human myeloid engraftment in the indicated organs after treatment completion (n=6/group) (n=3 independent experiments). **E.** *Ex vivo* clonogenic capacity of BM cells retrieved from mice treated as indicated (n=6/group). The left plot shows the number of colonies per 50,000 plated cells. The right plot, shows the total number of cells collected from each CFU plate. FISH analysis confirmed the leukemic MLL-AF9 identity of these cells. Percentages at the top of the FISH image indicate the percentage of MLLr+ cells detected in each indicated treatment (n=200 counted cells). Scale bar = 10µm. **F.** Scheme of the experimental design for secondary transplants. BM cells from treated primary mice were intratibially transplanted into secondary recipients at specific doses. Human engraftment was periodically monitored by PB and BM analysis. **G.** LSC estimation in secondary recipients using ELDA software. Mice were considered leukemic when presenting >0.1% human cells in BM (n=3 mice/dose and group). **H.** Human engraftment in BM at the end of the experiment for AraC- and combo-treated mice. **I.** Expression of the indicated HIF target genes identified in the scRNA-seq analysis to be overexpressed in the LSC^34^ cluster, in BM cells of mice treated with the indicated drugs (n=5-6 mice/group). WBC: white blood cells; RBC: red blood cells; BM: bone marrow; PB: peripheral blood; d: day. Data are shown as mean ± SEM. \**P* <0.05; ** *P* <0.01; Students’ t test analysis.

Limiting BM-derived AML cell doses from treated primografts were next serially transplanted into secondary recipients to further assess the impact of the AraC+BAY87 combo treatment for the long-term leukemia-initiating capacity of AML-LSCs (**Figure 7F**). A significant decrease in AML-LSC frequency (1/<44,063 *vs* 1/34,722, *P*=0.0272) was observed in secondary recipients transplanted with combo-treated *versus* AraC- treated primograft cells (**Figure 7G** and **S7D**), which was coupled to a 5-fold decrease in the leukemia burden 22 weeks after transplantation (**Figure 7H**). We then analyzed in the remaining resistant primograft cells the gene expression of HIF targets differentially expressed in the LSC^34^ cluster by scRNA-seq analysis (**Figure 7I**). Of note, we found a higher expression of *HIF1A*, *ENO1*, *PDK1*, *ALDOA1, NPM1, BNIP3 and ANXA1* in survivor BM cells from combo-treated mice than in AraC-treated counterparts, indicating an increased activation of the hypoxia pathway in chemotherapy-resistant cells. Collectively, these data support the *in vitro* results and indicate a synergistic effect of BAY87 with AraC treatment in debulking AML and eliminating AML-LSCs *in vivo*.

## Discussion

Although our understanding of the molecular and phenotypic features of AML is improving, yet many patients fail to respond to current treatments or exhibit early relapse. From the first reports of LSCs, leukemia ontogeny has been built upon paradigms of healthy hematopoiesis (3, 69). However, the classical view that LSCs are both rare and uniform, akin to normal HSCs, has gradually been revisited based on seminal studies performed in AML (69). Furthermore, studies investigating the biology of LSCs in AML use mainly murine models, and do not typically distinguish between the molecular subgroups used to stratify patients by risk when primary patient samples are used.

Here, we provide an exhaustive analysis at the single-cell level of the hypoxia/HIF signaling pathway in AML-LSCs in paired Dx-REL samples from pediatric/young adult risk-stratified AML patients. Owing to the great heterogeneity of AML disease and the complex functional interactions of different fusion proteins with HIFs, we focused on three specific cytogenetic subgroups. We resolved the intercellular transcriptional heterogeneity using scRNA-seq, which enabled us to identify and transcriptomically characterize the LSC population, providing to the best of our knowledge, the largest and most comprehensive single-cell expression atlas (119,000 cells) of AML cells and AML-LSCs to date.

We confirmed several features previously described for LSCs, including several LSC signatures, low ROS content, a more quiescence state, and a high activation of the translation process. These results are in accord with a recently published study analyzing 813 LSCs from 5 AML Dx-REL matched samples (70), and support clinical trials combining the proteasome inhibitor bortezomib to standard chemotherapy in AML (71). Strikingly, we consistently found an inverse correlation between the hypoxia signature and cell stemness, manifested as a gradual enrichment in hypoxia signature from LSC^34^ to differentiating NonLSC^34^ and NonLSC^38^ cells. This contrasts with earlier reports showing a higher activation of HIFs in the LSC population (13). Also, studies in healthy HSCs have shown the preferentially expression of *Hif-1a* in the more stem population (11) or in the more differentiated fraction (72) in BM cells of different mouse models. This incongruity might be explained by the high heterogeneity of AML patients analyzed in previous studies in absence of risk-stratification, the different phenotypic strategies to identify *bona fide* LSCs, or even by the use of distinct murine-based LSC readouts/approaches.

A hypoxia risk signature with prognostic value has been proposed (73), linking high HIF expression to shorter overall survival, similar to other studies (29, 72, 74). Comparison of paired Dx-REL samples enables the analysis of both therapy naïve- and therapy-resistant LSCs, providing insights into their evolution within the same patient. In this sense, our transcriptomic analysis revealed a patient-specific heterogeneity of both LSC6 and hypoxia scores in the seven paired Dx-REL samples. The relatively low number of patients included in the present study, however, does not allow us to draw clinico-biological conclusions.

Of note, and in line with other studies (29, 72), while LSC^34^ showed the lowest hypoxia enrichment score among the distinct analyzed clusters, it was nevertheless consistently enriched in both total AML cells and LSC^34^ cells when compared with both whole healthy BM cells and healthy CD34+ cells, regardless of the cytogenetic subgroup. Moreover, therapy (AraC)-resistant blasts have been reported to bind pimonidazole, an exogenous marker of hypoxia (65), encouraging us to explore the chemosensitizer role of HIFs inhibition in human AML. Indeed, targeting HIF1A has been explored as a therapeutic strategy in different malignancies (13, 19), and also its combination with AraC has also been tested in chronic lymphocytic leukemia (75) and in JAK2V617F-positive myeloproliferative neoplasms (76). We found a reduction in the LSC frequency *in vitro* when combining BAY87 and AraC. These results are in line with a previous report that tested the LSC dose in AML cells treated with AraC comparing normoxia and hypoxia culture conditions (77). We observed a similar chemoprotective effect of the low oxygen conditions when chemically manipulating the oxygen sensing ability of the cells. We also found a significant effect of the BAY87 and AraC combination *in vivo,* decreasing not only the presence of total AML cells but also of LSCs. We observed an increment in the LSC frequency in the AraC group with respect to control, consistent with a previous study describing an increment of CD34+ and progenitor cells after AraC treatment (60). This synergistic effect *in vivo* was, however, less dramatic than that observed *in vitro*. We speculate that the BM niche has a protective effect not present in the *in vitro* assays. Furthermore, optimization of the drug posology will be needed to completely unlock the potential of BAY87 as chemosensitizer.

In sum, we provide the largest and most comprehensive single-cell expression atlas (119,000 cells) of AML cells and AML-LSCs in paired Dx-REL samples from pediatric/young adult risk-stratified human AML patients to date. Our data indicate that the HIF/hypoxia pathway is attenuated in AML-LSC^34^ cells as compared with differentiated AML cells but it is enhanced when compared with healthy BM cells and HSPCs. Accordingly, chemical inhibition of the HIF pathway cooperates with standard-of-care chemotherapy to impair leukemogenesis *in vitro* and *in vivo*, substantially eliminating AML-LSCs. These findings support HIF pathway as a stem cell regulator in human AML and open new avenues for combinatorial targeted and chemotherapy-based treatments to specifically eliminate AML-LSCs.

## Acknowledgments

We thank Dr. Aleix Prat for technical help and Dr. Fernando Anjos-Afonso for technical discussions and advices. We thank the Finnish Hematology Registry and clinical Biobank (FHRB), Instituto Aragonés de Ciencias de la Salud (IACS) and the Blood Cancer UK Childhood Leukaemia Cell Bank for providing AML samples. We thank CERCA/Generalitat de Catalunya and Fundació Josep Carreras-Obra Social la Caixa for core support. Competitive financial support for this work was obtained from the Deutsche Josep Carreras Leukämie-Stiftung (DJCLS15R/2021) to PM and TV-H. This research was also supported by the Spanish Ministry of Economy and Competitiveness (SAF2016-80481R, PID2019-108160RB-I00), La Caixa Health Research Program (LCF/PR/HR19/52160011), the Leo Messi Foundation, “Heroes hasta la médula” initiative and ISCIII-RICORS within the Next Generation EU program (plan de recuperación, transformación y resiliencia) to PM and the Health Institute Carlos III (ISCIII/FEDER, PI20/00822) to CB. TV-H was supported by a Marie-Sklodowska Curie Fellowship (GA792923). JLT was supported by a Juan de la Cierva postdoctoral fellowship (FJC2019-040868-I). O.M. was supported by an investigator fellowship from the AECC (INVES211226MOLI).

## Author contributions

TV-H conceived the study, designed and performed experiments, analyzed and interpreted data, prepared figures and wrote the manuscript. JLT analyzed and interpreted scRNA-seq data, prepared figures and wrote the manuscript. AC and EE analyzed and interpreted bulk RNA-seq data. MV, FG-A, OM, VRC, PP, PR, RC, RDG and PL performed experiments. SB, LV and FA-J performed bioinformatic analyses. AB, HL, RWS, SV, MT and JLF provided human primary samples. CB and HH supported the study technically. PM, conceived the study, designed experiments, interpreted data, wrote the manuscript, and financially supported the study. All authors have read and agreed to publish the manuscript.

## Declaration of interests

PM is founder of the spin-off OneChain Immunotherapeutics which has no connection with the present research. The remaining authors declare no competing interests.

## Online Materials and Methods

### Analysis of public bulk-RNA-seq data

RNA-seq data from the publicly available databases TARGET (21), including a total of 78 patients in 147 RNA-seq runs, and Leucegene (22, 43), including a total of 72 patients in 302 RNA-seq runs, were downloaded for analysis. AML samples from specific cytogenetic subgroups without mutations in *TET2*, *IDH1* and *IDH2* were selected. **Table S1** summarizes the main clinico-biological features of the analyzed samples and the RNA-seq ID numbers. A total of 119 HIF target genes characterized by hypoxia-dependent transcriptional induction and the presence of functional hypoxia response elements were used to define the hypoxia transcriptomic signature (26) (**Table S2**).

#### Pre-processing and sample alignments

All samples were processed with the same pipeline and FastQC (78) was used for quality control and confirmation of the sequencing data from the FASTQ files. FASTQ files SRA for TARGET samples were extracted using the SRAToolKit (v 2.9.0) (https://github.com/ncbi/sra-tools).

#### Gene expression quantification

Illumina paired-end RNA-seq reads were aligned to the Gencode transcriptome release 27 (GRCH38.p10) (79) using Salmon (v0.7.2) (80). Quantification at gene level was performed using pseudo counts from Salmon quantification and transformation to counts per gene using *tximport* library function from Bioconductor (81).

#### Differential expression analysis

The following AML cytogenetic subgroups were included in the study: NK, inv(16), MLLr, t(8;21), FLT3^ITD^ and NPM1^mut^. The read counts per gene were transformed to log2 counts per million (logCPM) using edgeR (82) and those genes with mean logCPM < 0 were filtered out. Normalization of the data was performed using the TMM method from edgeR package. Differential expression analysis was performed with LIMMA (83) using the function limma.voom adjusted by SVA (84).

#### Functional enrichment analysis

GSEA was conducted (85) based on the hypoxia transcriptomic signature described above, using the pre-ranked enrichment method, sorting all the genes by –*log*_10_ (*p* – *value*) obtained from the differential expression analysis.

### Primary AML cells

Primary AML samples were obtained from accredited Biobanks (Finnish Hematology Registry and clinical Biobank (FHRB), Instituto Aragonés de Ciencias de la Salud (IACS) and the Blood Cancer UK Childhood Leukaemia Cell Bank) and from collaborating hospitals (Hospital Clinic of Barcelona, Barcelona, Spain; Hospital Princess Maxima, Utrech, The Netherlands; Hospital Germans Trias i Pujol, Badalona, Spain; Hospital Sant Joan de Deu, Barcelona, Spain; and Hôspital d’enfants Armand Trousseau, Paris, France). Samples were obtained from routine diagnostic procedures after written consent from patients or parents/guardians in case of minors. The study was approved by the Institutional Ethical Review Board of Hospital Clinic of Barcelona (HCB/2018/0020). AML mononuclear cells were frozen until use in liquid nitrogen using fetal bovine serum (FBS) (Sigma) with 10% dimetylsulfoxide (Sigma). The mutational state of AML samples was analyzed on DNA extracted from total cells using the Maxwell RSC Blood DNA Kit (Promega) and a next generation sequencing (NGS) panel of mutations using the Oncomine Myeloid Research Assay (ThermoFisher). **Table S3** lists the main clinico-biological features of the AML samples used in this study.

### Single-cell RNA sequencing

### Sample preparation

Frozen BM AML cells were thawed and stained (30 min at 4°C) in PBS + 2% FBS with the following antibodies: anti-hCD45-BV510 (HI30), anti-hCD33-BV421 (WM53), anti-hCD34-APC (581) and anti-hCD38-FITC (HIT2) (all from BD Biosciences). Cells were washed with PBS, filtered through a 40-µm strainer and stained with 7AAD (1:100, BD Pharmingen) for 5 min before sorting in FACS Aria-II Fusion cell sorter (BD Bioscience) using a 100-µm nozzle. A minimum of 20,000 cells of each CD34+CD38- (LSC-enriched population) and CD34-CD38+ (LSC-depleted population) sample were collected in PBS + 2% FBS for downstream applications.

#### Library preparation and sequencing

The cell number and viability of the CD34+CD38- and CD34-CD38+ samples were verified with a TC20™ Automated Cell Counter (BioRad Laboratories) and cells were partitioned into Gel Bead-In-Emulsions using the Chromium Controller system (10X Genomics), with a target recovery of 5,000 total cells of each population. cDNA sequencing libraries were prepared using the Next GEM Single Cell 3’ Reagent Kit v3.1 (10X Genomics, PN-1000268). Briefly, after GEM-RT clean up, cDNA was amplified during 12 cycles and cDNA QC and quantification were performed on an Agilent Bioanalyzer High Sensitivity chip (Agilent Technologies). cDNA libraries were indexed by PCR using the PN-220103 Chromiumi7 Sample Index Plate. Size distribution and concentration of 3’ cDNA libraries were verified on an Agilent Bioanalyzer High Sensitivity chip (Agilent Technologies). Finally, sequencing of cDNA libraries was done on the Illumina NovaSeq 6000 platform using the following sequencing conditions: 28 bp (Read 1) + 8 bp (i7 index) + 0 bp (i5 index) + 89 bp (Read 2), to obtain approximately 25-30,000 reads per cell.

#### scRNA-seq data analysis

Reads were aligned to the Hg38 *Homo sapiens* reference genome and quantified through CellRanger Single-Cell Software Suite (v3.1.0). Each sample was analyzed individually prior to data integration. Low-quality cells were filtered out based on mitochondrial RNA percentage, number of unique molecular identifiers (UMIs), and number of different genes (thresholds adjusted separately for each data set). The CD34+CD38- and CD34-CD38+ libraries were merged for each sample before applying usual processing following Seurat tutorials (highly variable genes calculation, log-normalization, scaling and correction by number of UMIs and mitochondrial content). Seurat v4.0.1 was used (86) for R 3.6.1. Principal component analysis (PCA) was performed with a number of principal components ranged between 10 and 20, depending on data set complexity. Dimensionality reduction was performed by applying Uniform Manifold Approximation and Projection (UMAP) algorithm.

The selection of LSC clusters was done independently on each sample. We assigned an LSC6 score for each cell using the six gene signature and weights proposed in Elsayed *et al*, 2020 (34). Due to the sparse nature of the single-cell data, rather than selecting the cells with highest LSC6 score, we elected to cluster the data in an unsupervised manner using the Louvain clustering algorithm with resolution values ranging from 0.5 to 1, and rank the obtained partitions according to their average LSC6 score. Those clusters above LSC6 decile 9 were determined as the more likely to be enriched on LSCs. If more than one cluster was selected under these criteria, the proportions of *in-silico* predictions obtained from VanGalen *et al*, 2019 (36) and Triana *et al*, 2021 (37) were used. The cluster with the highest enrichment of HSC-like predicted cells was finally determined as the most likely to be enriched on LSCs. Cell cycle phases identification was performed based on previously defined markers (87). Scripts and plots generated on each sample are available in Github (https://github.com/JLTrincado/scAML).

#### *In-silico* prediction of cell types

Some studies have reported phenotypic heterogeneity in human BM. We leveraged these annotated datasets to predict the healthy cell type closest to our leukemic clusters. The annotated healthy BM datasets from Van Galen *et al* (36) was merged and projected onto each sample using FindTransferAnchors and TransferData methods from Seurat (86). Code for reference assembly and projection is available at Github (https://github.com/JLTrincado/scAML). For projecting the data onto healthy BM data from Triana *et al* (37), a workflow based on scmap (88) was used. Sample code for reference atlas projection is available at https://git.embl.de/triana/nrn//tree/master/Projection_ Vignette.

#### Integration by cytogenetic-molecular subgroup

Seurat canonical correlation analysis (CCA, number of anchors set to 2,000) was applied to correct the patient-specific bias introduced by the pooled transcriptomic information from all sequenced samples (86). Individual clusters identified in each sample to be enriched in LSCs, were labeled in the integrated datasets as “LSC^34^”. All the remaining cells non-labeled as “LSC^34^” within the CD34+CD38- population were labeled as “NonLSC^34^”. All CD34-CD38+ cells were labeled as “NonLSC^38^”.

#### Pathway scores and pseudotime trajectories

Different gene sets reported in the literature to be associated with LSC-enriched pathways (**Table S2**) were used to biologically inspect each annotated cluster. AddModuleScore from Seurat suite was used to assign a score to each cell for each gene set (86). Resulting values were normalized between 0 and 1. Trajectory analyses were performed with the Monocle package (v2.18.0) (89). The highly variable genes obtained for the integration of the data via Seurat were used for pseudotime ordering. Dimensionality reduction was applied with the DDRTree option.

### Cell lines

THP-1, Kasumi-1, ME-1 and MS5 cell lines were purchased from the DSMZ German Collection of Microorganisms and Cell Cultures (Braunschweig, Germany). THP-1, Kasumi-1 and ME-1 were cultured in RPMI-1640 supplemented with GlutaMAX (Gibco), 10% FBS (20% for ME-1) and penicillin-streptomycin (P/S) (Gibco) at 37°C with 5% CO_2_. MS5 cells were cultured in αMEM (Gibco) with 10% FBS and P/S. MS5 cells were irradiated (50 Gy) and seeded on collagen (StemCell Technologies)-coated plates as monolayers for co-culture with primary AML cells. Cells were passaged every 2-3 days and maintained in an exponential growth phase. All cultures were routinely tested for mycoplasma.

### Xenotransplantation

Eight- to twelve-week-old non-obese diabetic (NOD).Cg-*Prkdc^scid^Il2rg^tm1Wjl^*/SzJ (NSG) mice (The Jackson Laboratory) were bred and housed under pathogen-free conditions. The Animal Care Committee of the Barcelona Biomedical Research Park approved all experimental procedures with mice (HRH-17-0014 and HRH-19-0003). A total of 0.3–1 × 10^6^ primary AML cells were intra-BM transplanted into sublethally irradiated (2 Gy) NSG mice (90). AML cells were previously incubated 30 min at 4°C with OKT3 (BioXCell). Human engraftment was monitored through PB and BM aspirates from week six after transplantation until AML graft levels were ∼20% in BM or ∼2% in PB. Mice were then homogeneously divided into the following treatment groups (n=5-6/group): (i) AraC (and carrier solution), (ii) BAY 87-2243 (and PBS), (iii) AraC and BAY 87-2243, and (iv) control (PBS and carrier solution). Cytarabine/AraC (50 mg/Kg, Accord) was administered intraperitoneally for 5 days (60, 65). BAY 87-2243 (4 mg/Kg, Selleckchem) was administered for 5 days by oral gavage (64). Mice were sacrificed 48-72 h after treatment completion and PB, BM, spleen and liver were collected to analyze the efficacy of each treatment. White and red blood cell and platelet counts were determined with a hemocytometer (2800VET V-Sight, Menarini Diagnostics). To assess the frequency of AML-LSCs, BM-derived mononuclear cells were collected from primografts (two different mice with similar human engraftment per treatment group) and were intra-BM transplanted into irradiated (2 Gy) secondary NSG recipients (n=5/group/cell dose) and were analyzed as above.

### Immunophenotyping and cell cycle, apoptosis and CellROX analyses

#### Immunophenotyping

AML engraftment in mice was monitored by FACS analysis, biweekly in PB and at sacrifice in PB, BM, spleen and liver. PB was collected into EDTA tubes (Sarstedt). Mononuclear cells were stained (30 min at 4°C) with the following monoclonal antibodies: anti-hHLA-ABC-FITC (G46-2.6), anti- hCD45-APC (HI30), anti-hCD33-PE (WM53), anti-hCD34-PECy7 (8G12) and anti-hCD19-BV421 (HIB19) (all from BD Biosciences). Cells were then lysed and fixed with the BD FACS^TM^ Lysing solution (BD Biosciences). Fluorescence Minus One (FMO) controls were used to set the FACS gates. A FACSCanto^TM^- II flow cytometer and equipped with FACSDiva^TM^ software was used for analysis (BD Biosciences).

#### Cell cycle analysis

Cells were stained with anti-hCD45-BV510 and anti-hCD33-BV421 for 30 min at 4°C.

After washing, cells were fixed with 0.4% paraformaldehyde (Alfa Aesar) for 30 min at room temperature (RT), then lysed with 0.2% TritonX (Sigma) for 1 h at 4°C, washed, stained with anti-Ki67-PE (1:20, BD Biosciences) for 2 h at 4°C and finally stained with 7AAD (BD Bioscience) for an additional one hour. Cells were analyzed using a FACSCanto^TM^-II flow cytometer and equipped with FACSDiva^TM^ software.

#### Apoptosis

Cells were washed with Binding Buffer 1X (BD Pharmingen) and stained with anti-hCD33-BV421, anti-hCD45-BV510, anti-hCD34-APC and anti-hCD38-FITC for 30 min at 4°C. Cells were then washed with Binding Buffer 1X and stained with AnnexinV-PE (BD Biosciences) and 7AAD for 15 min at RT. Cells were analyzed within an hour using a FACSCanto^TM^-II flow cytometer and equipped with FACSDiva^TM^ software.

#### CellROX

For ROS content analysis, cells were stained with anti-hCD33-BV421, anti-hCD45-BV510, anti-hCD34-PE (581), anti-hCD38-FITC and with CellROX Deep Red Reagent (1:500, ThermoFisher) for 30 min at 37°C. Cells were washed 3 times with PBS and analyzed using a FACSCanto^TM^-II flow cytometer and equipped with FACSDiva^TM^ software.

### Clonogenicity and LTC-IC assays

The clonogenic capacity of leukemic progenitors was evaluated in CFU assays. AML cells (500-50,000 cells/well) were seeded in semisolid methylcellulose media (MethoCult #H4434; StemCell Technologies) according to manufactureŕs instructions. Triplicates of each sample/primograft were seeded. CFU numbers from primograft AML cells were normalized to the total human engraftment of each particular donor mouse.

LTC-ICs assays were conducted to evaluate the LSC frequency after *in vitro* treatment with drugs (33, 77). In brief, primary AML BM samples were thawed and seeded on confluent MS5 monolayers on MyeloCult H5100 (StemCell Technologies) supplemented with human IL3 (Miltenyi Biotec), human G-CSF (Amgen) and human TPO (PeproTech) at 20 ng/mL each and 1X P/S (Gibco). Cells were allowed to recover for 48 h and were then treated with the corresponding drugs and maintained for 48 h at 5% O_2_ (hypoxic conditions). After drug treatment, AML-MS5 co-cultures were harvested and MS5 cells and T cells were magnetically depleted by AutoMACs (Miltenyi Biotec) using anti-murine Sca1 and anti-human CD3 magnetic beads (Miltenyi Biotec). Recovered cells were counted and different doses (2,000, 1,000, 500 and 250 cells) were seeded each in 15 wells of a 96-well plate pre-coated with MS5 cells in supplemented MyeloCult media and allowed to expand in 20% O_2_ (normoxic conditions) with media changes twice weekly. After 5 weeks, wells were score as positive if massive growth of cells were observed in the well (33). LSC dose was determined using ELDA software (91). The identity of the AML cells was confirmed by detection of the molecular alteration by FISH or qPCR in some of the positive wells.

### Fluorescence in situ hybridization (FISH)

Cells were resuspended in hypotonic solution (0.075 mM KCl) for 20 min at 37°C and fixed in cold methanol:acetic acid (3:1). Samples were spread onto methanol-cleaned slides and kept at -20°C until processing. Two-color FISH experiments were performed using either XL CBFB, XL t(8;21) (both from MetaSystems) or LSI MLL Break-Apart (Abbott Molecular) probes to detect inv(16), t(8;21) or MLL rearrangements, respectively. FISH was performed following standard procedures (90, 92, 93). Briefly, cells were denatured at 73°C in 70% formamide in 2×SSC for 2 min. Hybridization was carried out by adding 5 μl of the DNA probe mixture to preparations and incubating the slides in a humid chamber at 37°C for 16 h. Post-hybridization washes were performed in 0.4×SSC with 0.3% NP-40 at 73°C followed by 2×SSC with 0.1% NP-40 at RT, for 1 min each. Slides were mounted with DAPI II solution (Abbott Molecular). Analyses were performed using a Nikon Ci-S/Ci-L epifluorescence microscope equipped with specific filters for DAPI, FITC, Cy3 and a dual-band pass filter for FITC and Cy3. A minimum of 200 informative nuclei were analyzed per experiment.

### RNA purification and gene expression profiling

RNA was extracted from a pellet of 0.5-1 x 10^6^ cells using a Maxwell RSC simply RNA Cells Kit (Promega) on a Maxwell RSC system (Promega). Between 0.2-2 µg of RNA were reverse-transcribed into cDNA using the SuperScript III Reverse Transcriptase (Invitrogen) following manufacturer’s instructions. cDNA samples were used as templates for real-time PCR analysis using SYBR Green Mastermix (Invitrogen) on a BIORAD CFXTM Real-Time system (Bio-Rad). Oligonucleotides used are detailed in **Table S4**. Gene expression was normalized with respect to the expression to the housekeeping gene *GUSB*.

### Drugs

AraC and BAY 87-2243 were purchased from Accord and Selleckchem, respectively. AraC was used at 3 µM *in vitro* and at 50 mg/kg/body weight *in vivo*, administered intraperitoneally daily for 5 days, as described (65). Control animals were treated with the same volume of PBS. BAY 87-2243 was used at a final concentration of 10 mM *in vitro,* previously dissolved in ethanol (Scharlau). Control cells were treated with same amount of ethanol. For *in vivo* experiments, BAY 87-2243 was dissolved in carrier solution (10% ethanol, 40% solutol HS15 (Sigma), 50% sterile distilled water) and administered orally by gavage (4 mg/kg/body weight) daily for 5 days, as previously described (64). Control animals were treated with the carrier solution.

### Statistical analysis

Data are represented as mean ± standard error (SEM). Statistical comparisons between groups were assessed using two-tailed unpaired Student’s t-tests, or paired Student’s t-tests (when analysing data from same AML samples subjected to different treatments), unless otherwise stated. Data distribution was assumed to normal but this was not formally tested. All analyses were performed with Prism software, version 8.0 (GraphPad software Inc., San Diego, CA) and *P*<0.05 was considered statistically significant (\**P*<0.05 and \*\**P*<0.01).

### Data and code availability

Newly generated scRNA-seq data have been deposited on the European Genome-Phenome Archive (EGA) and are accessible through accession no. EGAS00001005980. All analyses and code used along this study are available at https://github.com/JLTrincado/scAML. All other supporting data/reagents are available upon reasonable request.

## Supplemental Figures

**Figure S1.**
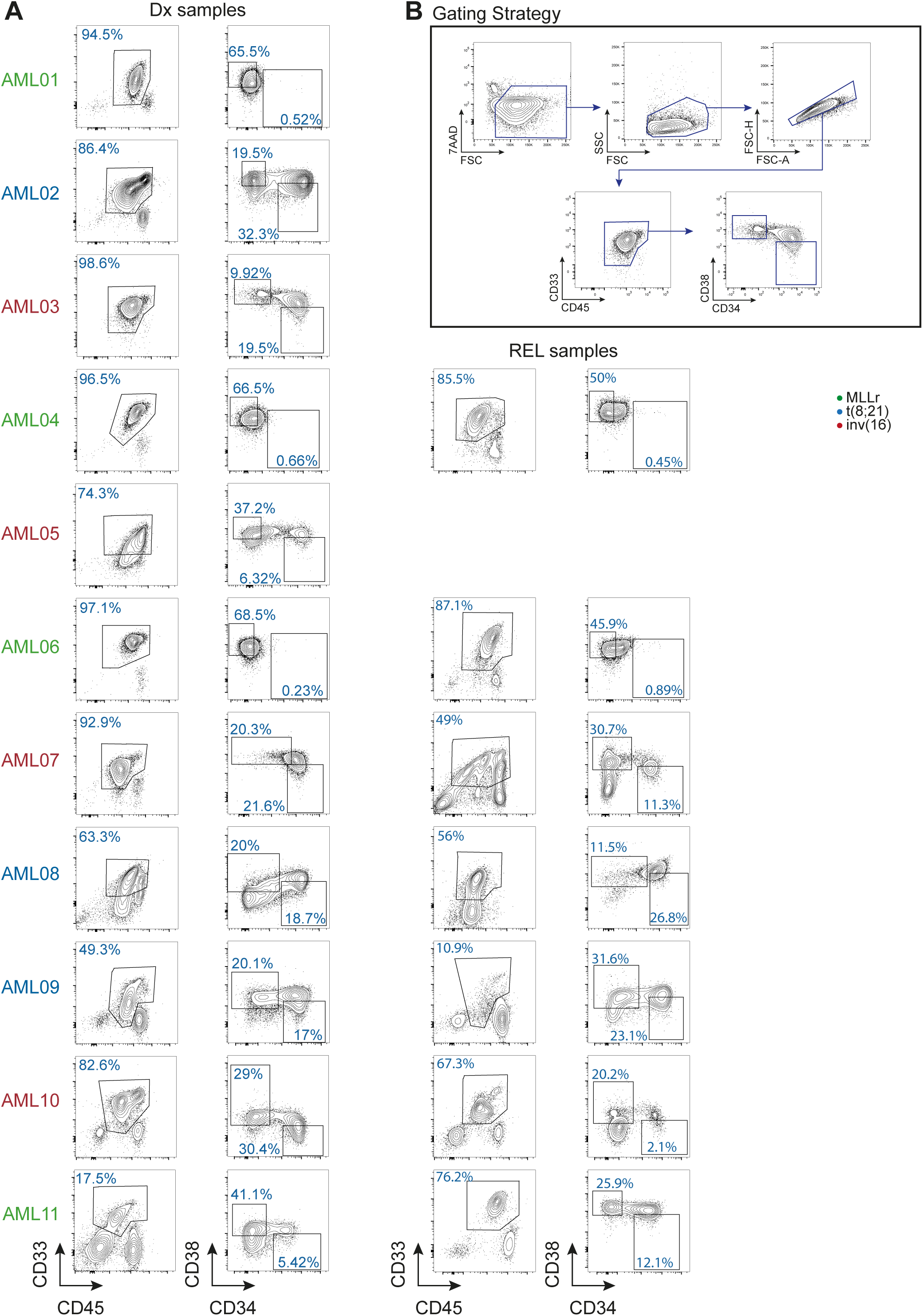
FACS analysis and sorting strategy for each AML sample used in this study (related to Figure 2). **A.** FACS plots showing the expression of CD45, CD33, CD34 and CD38 of each Dx and REL AML samples. **B.** Stepwise gating strategy used for FACS sorting of the CD34+CD38- and CD34-CD38+ AML subpopulations.

**Figure S2.**
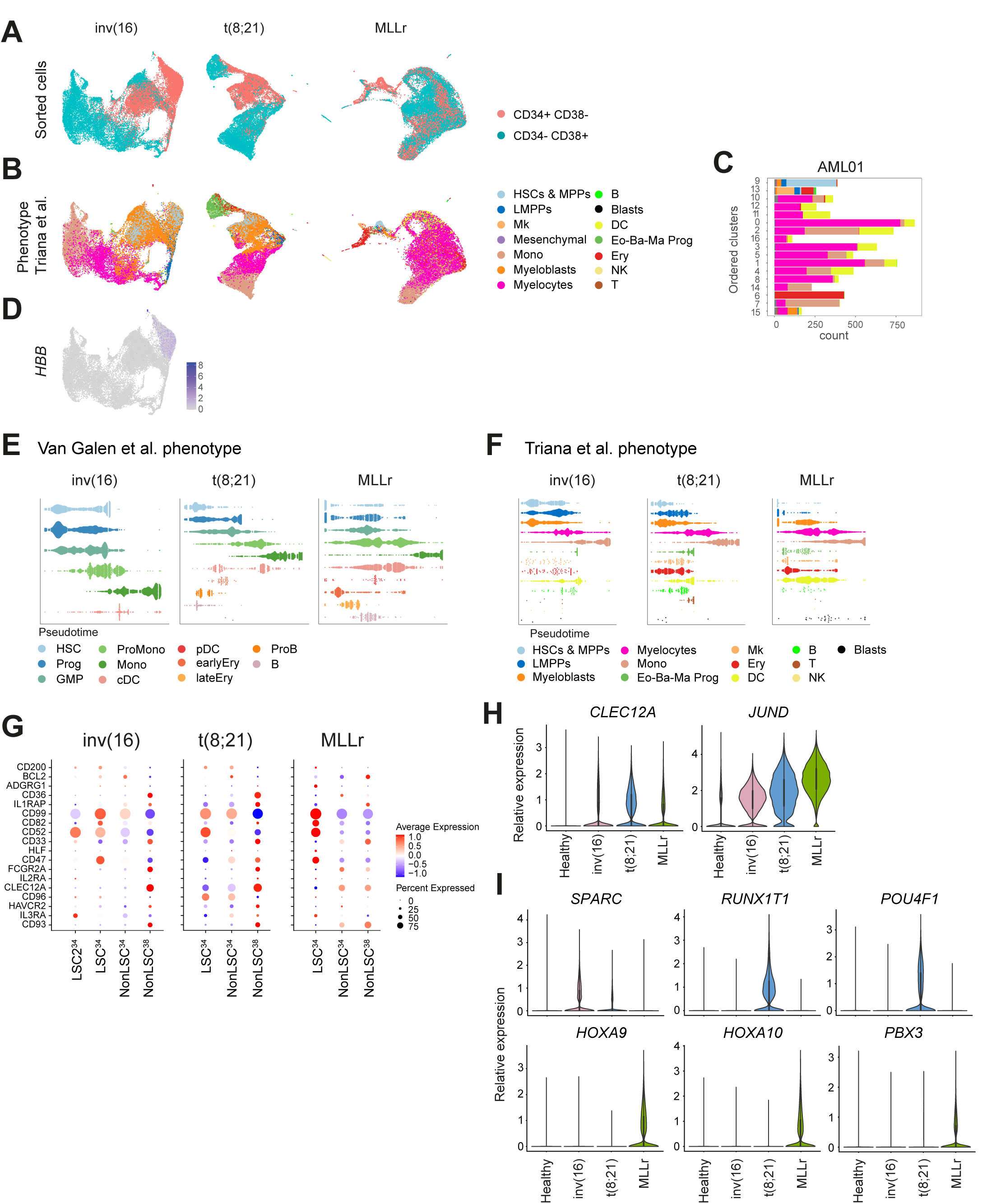
Single cell transcriptomic characterization of the sequenced AML cells (related to Figure 3). **A.** UMAP plots showing from which sorted population (CD34+CD38- or CD34-CD38+) each cell belongs integrating the samples from each cytogenetic subgroup. **B.** UMAP plots showing the predicted phenotype of the cells according to Triana *et at* for all the cells integrated from the different samples in each cytogenetic subgroup. **C.** Number of cells from each predicted phenotype according to Triana *et al* included in each cluster of sample AML01. **D.** UMAP plot showing the expression of *HBB* in the integrated inv(16) AMLs. **E-F.** Trajectory/Pseudotime analysis of the cells included in each of the defined phenotypes according to Van Galen *et al* (**E**) and Triana *et al* (**F**). **G.** Comparative relative expression of established stem cell markers in the different defined populations of AML cells. **H.** Expression of the AML markers *CLEC12A* and *JUND* in the different AML cytogenetic subgroups compared with healthy BM cells. **I.** Expression of the indicated genes in the different AML cytogenetic subgroups compared with healthy BM cells. Overexpression of *SPARC*; *RUNX1T1* and *POU4F1;* and *HOXA9*, *HOXA10* and *PBX3* is well-reported for inv(16), t(8;21) and MLLr AMLs, respectively. LSC: leukemic stem cell; HSC: hematopoietic stem cell; Prog: progenitor; GMP: granulocyte-macrophage progenitor; ProMono: promonocyte; Mono: monocyte; cDC: conventional dendritic cells; pDC: plasmacytoid dendritic cells; Ery: erythroid progenitor; ProB; B cell progenitor; B: mature B cell; Plasma: plasma cell; T: naïve T cell; CTL: cytotoxic T lymphocyte; NK: natural killer cell; Mk: megakaryocyte ; LMPPs: lymphoid primed multipotent progenitor; MPPs: multipotent progenitor; Eo-Ba-Ma Prog: eosinophil-basophil-mast cell progenitor.

**Figure S3.**
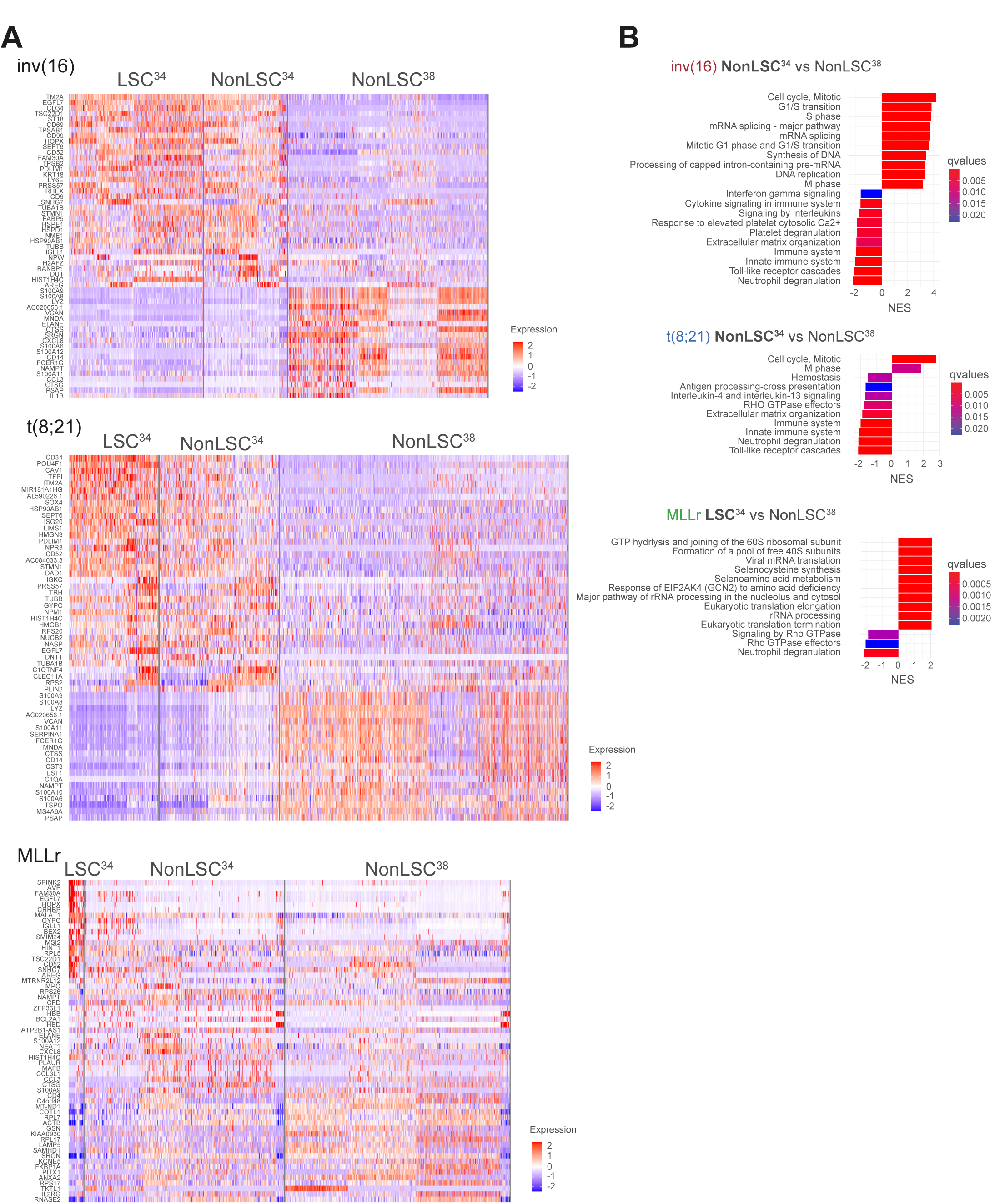
Differential gene expression analysis in the defined AML clusters (related to Figure 3). **A.** Heatmaps of the DEGs of each of the defined clusters in the 3 cytogenetic groups. **B.** GSEA showing the enriched pathways in the different defined clusters of AML cells. For inv(16) and t(8;21) AMLs comparison is shown between NonLSC^34^ and NonLSC^38^ clusters. For MLLr AMLs, comparison is made between LSC^34^ and NonLSC^38^ clusters.

**Figure S4.**
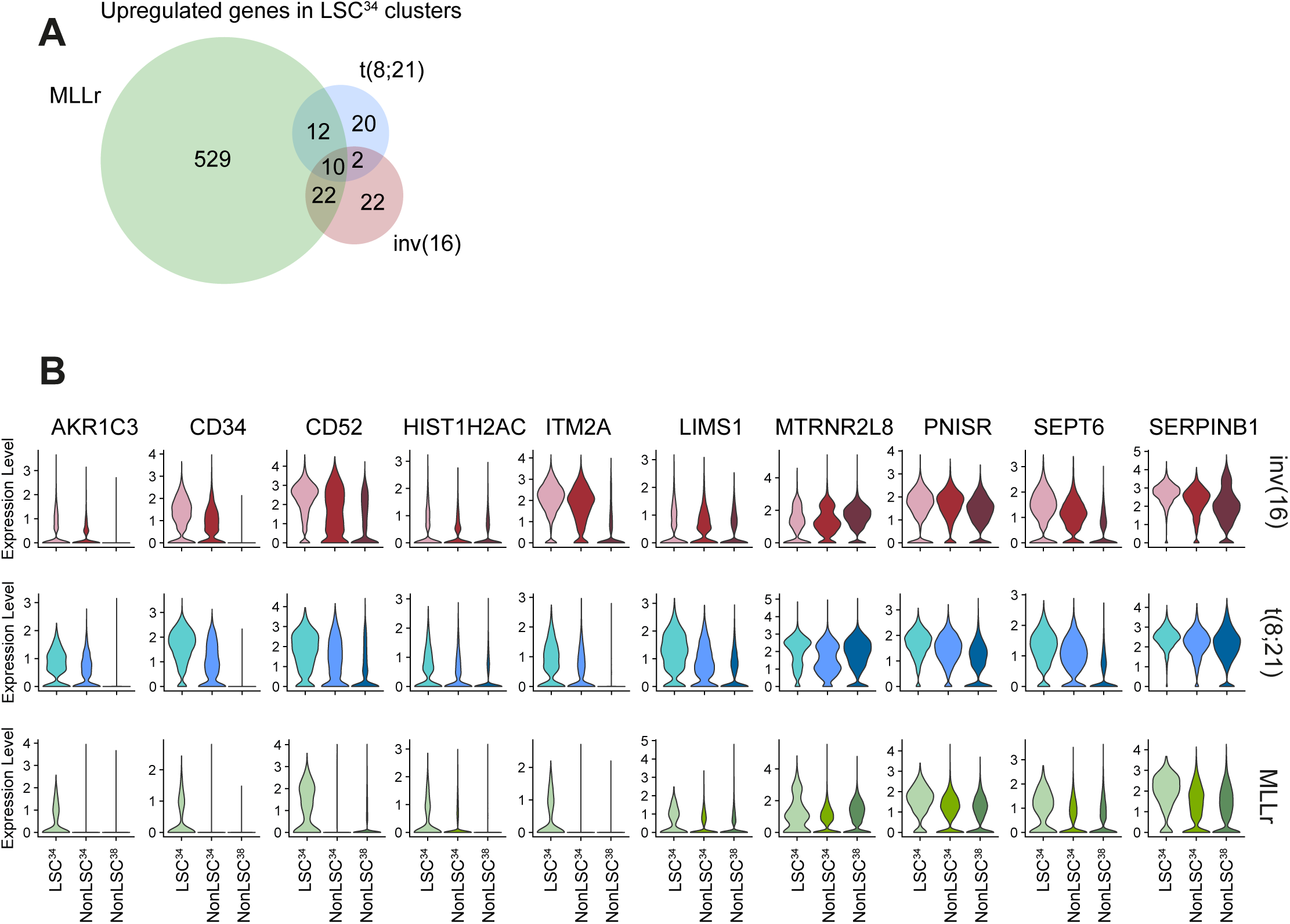
Upregulated genes in the LSC^34^ cluster (related to Figure 3). **A.** Venn diagram showing the number of significantly upregulated genes in the LSC^34^ cluster in the different cytogenetic AML subgroups. The number of upregulated genes shared by LSC^34^ cluster of distinct cytogenetic subgroups is also shown. **B.** Expression of the shared 10 genes specifically upregulated in the LSC^34^ clusters of the 3 distinct cytogenetic subgroups.

**Figure S5.**
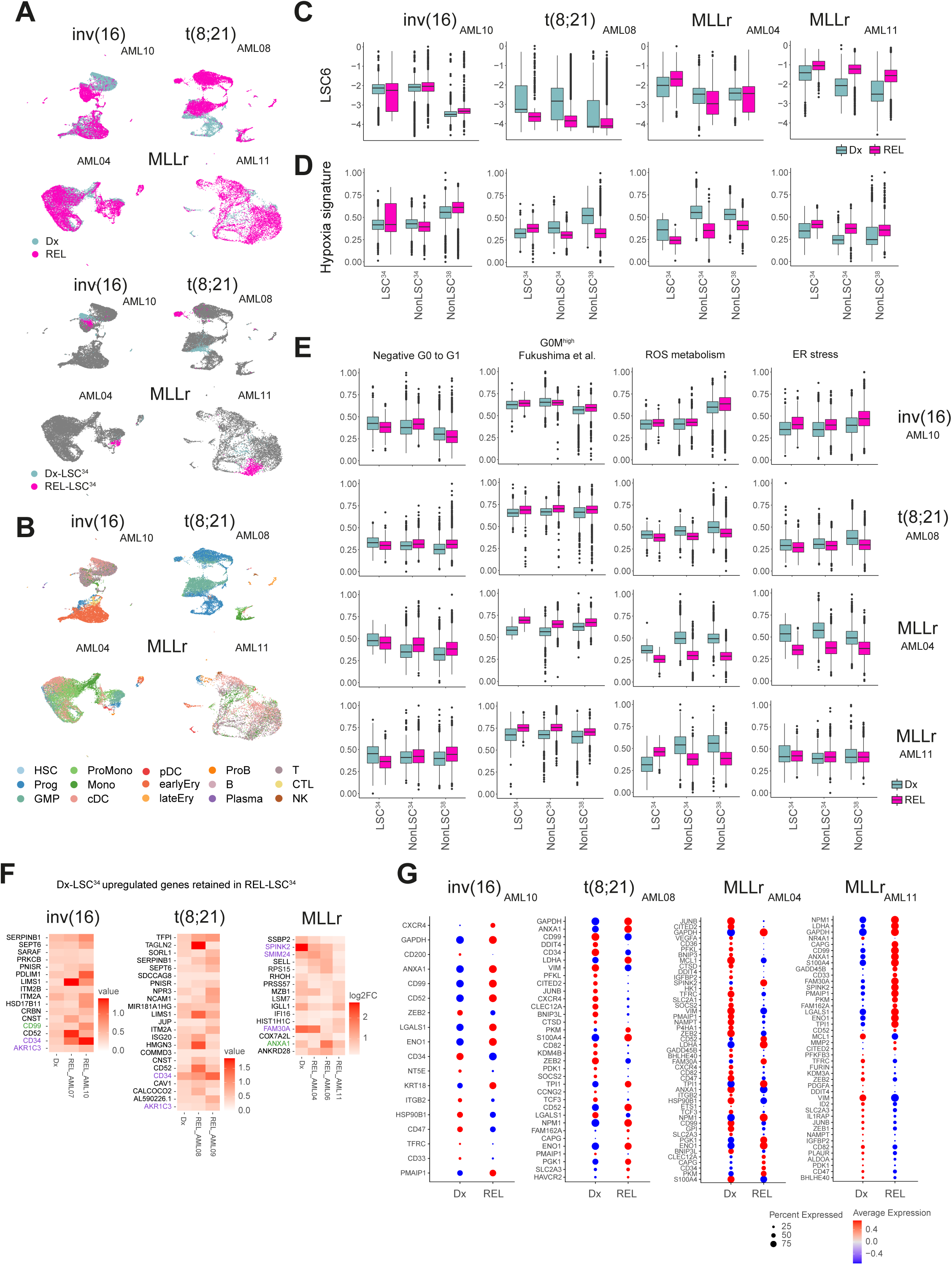
Single cell transcriptomics on paired Dx-REL samples (related to Figure 5). **A.** UMAP plots integrating Dx and REL AML cells from the indicated patients (top plots) and showing the identified LSC^34^ cluster at Dx and REL (bottom plots). **B.** UMAP plots showing the predicted phenotype according to Van Galen *et al* in the Dx and REL integrated AML cells from the indicated patients. **C-D.** LSC6 (**C**) and hypoxia (**D**) signature scores of the defined clusters in Dx and REL AML cells from the indicated patients/cytogenetic subgroups. **E.** Analysis of different metabolic pathways related to stemness and hypoxia in the defined clusters in Dx and REL AML cells from the indicated patients/cytogenetic subgroups. **F.** Genes commonly upregulated in the LSC^34^ clusters at both Dx and REL. In purple, genes included in the LSC6 score; in green, hypoxia target genes. **G.** Hypoxia target genes differentially expressed between Dx and REL in the indicated paired samples. HSC: hematopoietic stem cell; Prog: progenitor; GMP: granulocyte-macrophage progenitor; ProMono: promonocyte; Mono: monocyte; cDC: conventional dendritic cells; pDC: plasmacytoid dendritic cells; Ery: erytroid progenitor; ProB; B cell progenitor; B: mature B cell; Plasma: plasma cell; T: naïve T cell; CTL: cytotoxic T lymphocyte; NK: natural killer cell; LSC: leukemic stem cell; log2FC: log2 fold change.

**Figure S6.**
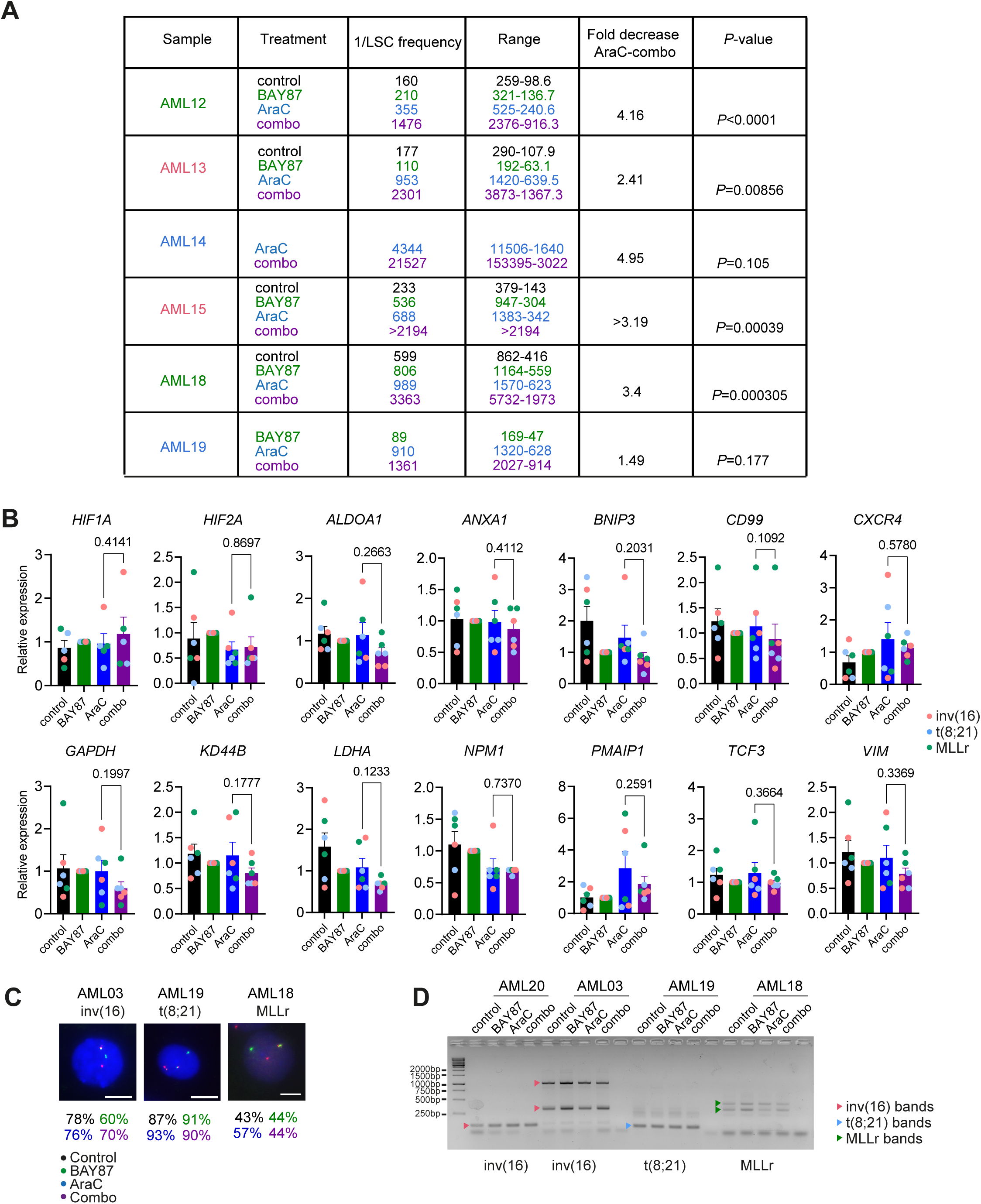
Inhibition of HIF pathway sensitizes AML-LSCs to chemotherapy (related to Figure 6). **A.** Detailed estimation of the LSC frequency at the completion of the LTC-IC assay with the ELDA software showing the complete results and differences among the AraC- and combo-treated cultures. **B.** Expression of the indicated HIF target genes (identified in the scRNA-seq analysis to be overexpressed in the LSC^34^ cluster) after 48 h of the indicated treatments at 5% O_2_ (n=6 samples, AML03, AML16-AML21). Statistical significance was calculated using the paired Students’ t test. Expression is normalized respect to the BAY87 samples. **C.** FISH analysis of the AML cells after 48 h treatment at 5% O_2_. Data indicate the percentage of cells harboring the AML-specific rearrangements inv(16), t(8;21) and MLLr. n=200 counted cells. Scale bar = 10µm. **D.** qPCR analysis of the treated AML cells, confirming the expression of the gene rearrangement transcript.

**Figure S7.**
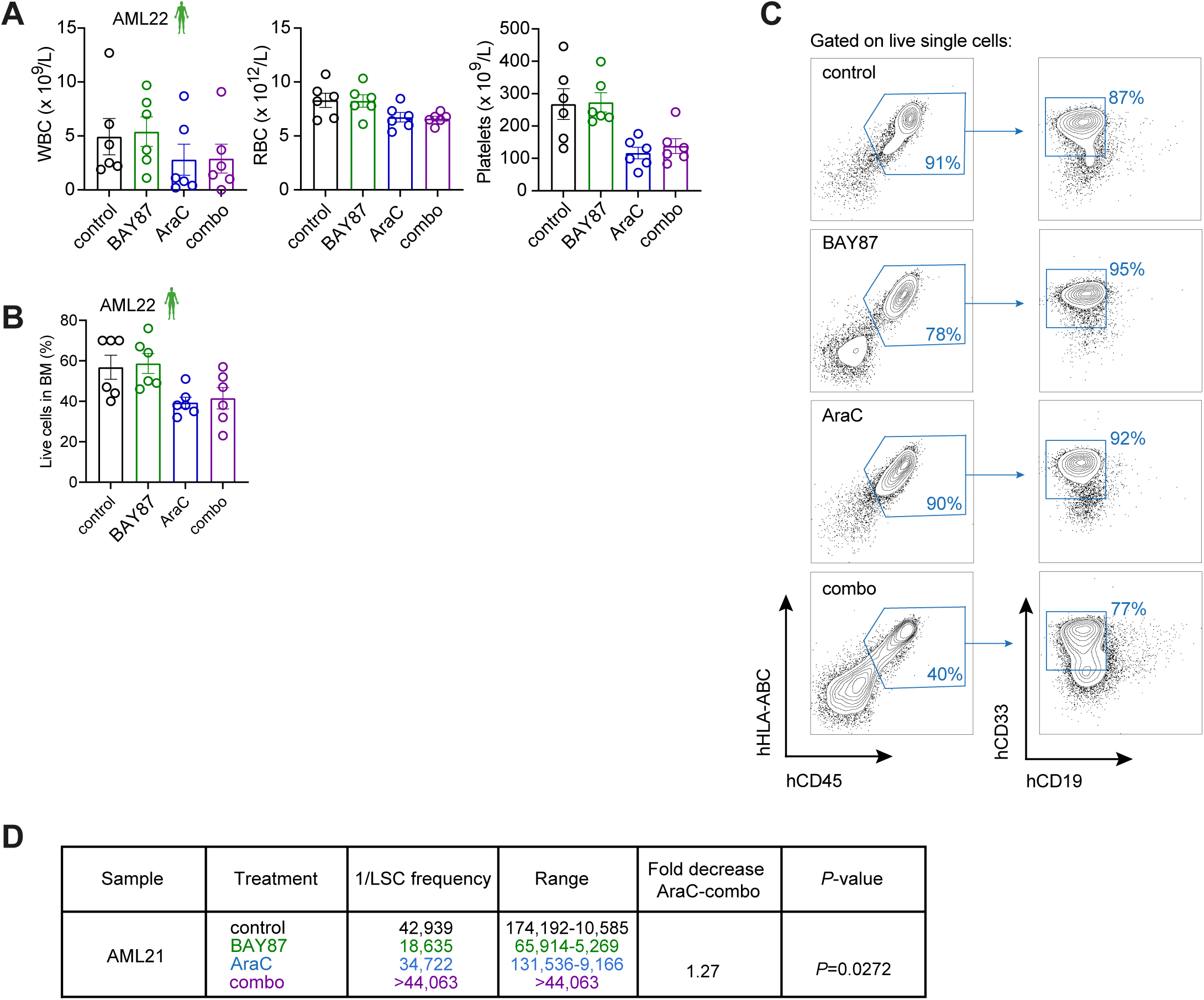
Inhibition of HIF pathway sensitizes AML-LSCs to chemotherapy *in vivo* (related to Figure 7). **A.** WBC, RBC and PLT counts in PB of mice treated as indicated (n=6/group). Representative data from one experiment (n=3). **B.** Total (mouse and human) BM live cells evaluated by trypan blue exclusion, in mice treated as indicated (n=6/group). **C.** Representative FACS plots of BM cells after completion of the treatment. Human myeloid (AML) engraftment was identified as hHLA-ABC+ hCD45+ hCD33+ hCD19-. **D.** Detailed estimation using the ELDA software of the LSC frequency (sample AML21) at the completion of the secondary transplants, reflecting the decrease of LSC dose in combo-treated AML xenografts.

Table S1. TARGET and Leucegene samples analyzed by bulk RNA-seq (related to Figure 1).

Table S2. Gene signatures (related to Figures 1-5).

Table S3. Primary AML samples used in this study (related to Figures 2-7).

Table S4. Primers used for qPCR (related to Figures 6-7).

## Notes

https://github.com/JLTrincado/scAML

### Competing Interest Statement

The authors have declared no competing interest.

